# Impaired axon regeneration and heightened synaptic dynamics in the injured aged mammalian cortex

**DOI:** 10.1101/2024.12.27.630526

**Authors:** Cher Bass, Anil A Bharath, Vincenzo De Paola

## Abstract

How aging affects axon regeneration and synaptic repair in the brain is poorly understood. To study age-related changes in neural circuits, we developed a model of axonal injury in the aged (*>* 2 years) mouse somatosensory cortex. By directly tracking fluorescently labelled injured axons by multiphoton imaging, we find that while axon degeneration in the aged brain is comparable to the young adult brain, axon regeneration is impaired. We further examine changes in the most common type of cortical synapses, En Passant Boutons (EPBs), and observe a transient and significant increase in the number and size of boutons 6 hours post-lesion. Using a computational model of a recurrent neural network, we examined the functional consequences of these synaptic increases on memory, comparing results to models of the young adult brain. The results suggest that increased synaptic dynamics might enable partial recovery from injury via synaptic re-wiring in the aged brain.

**Highlights:**

- Impaired axon regeneration in the aged brain is not due to slowed degeneration.
- Synaptic reorganization occurs following axon injury in the aged brain.
- Synaptic reorganization may serve as a compensatory mechanism after injury.

## Introduction

Aging is accompanied by impaired brain functions, including decline in sensory abilities^1,2^, reduced cognitive function^3–5^ and increased vulnerability to injury or disease^6–8^, but the underlying mechanisms are not well defined. I n o u r previous work, we compared young adult (2-3 months old) and aged mice (*>* 2 years old) and found an increase in axonal bouton (pre-synaptic terminals) dynamics in a subset of cortical neurons.^9^ En Passant Bouton (EPB)-rich axons which originate in Layer (L) 2/3/5, or the thalamus, rather than Terminaux Bouton (TB)-rich axons which originate in L6, increased their turnover rate (TOR; i.e. their formation and elimination), and rate of size change with age^9^. Thus, ageing may impact cognitive impairment^5^ by affecting synaptic re-wiring.^9,10^ In addition, ageing has an established detrimental effect on nerve recovery after injury in the spinal cord^6,11^, but its effect in the injured brain is less clear. Previous studies investigated how injury affects cortical axons in the young adult brain, and found a cell-type specific response where EPB-rich axons regenerated less (20%) than TB-rich axons (55%).^12,13^ Axon degeneration and regeneration in the brain are challenging to study as many current techniques rely on fixed tissue analysis, which provides only a snapshot of degeneration and requires large number of animals. Real-time, dynamic imaging is needed to understand the sequence of degeneration and repair processes in the brain.^14^

Here, we investigated how axonal injury affects axonal and synaptic responses in the aged somatosensory cortex with longitudinal *in vivo* multiphoton microscopy. We used laser-mediated axotomy, which is known to cause a small local lesion inducing only a transient recruitment of microglia and little to no formation of glial scar.^12,13,15^ While this lesion paradigm does not replicate the complexity of injury processes in diseases such as traumatic brain injury^16^, it provides a valuable model for studying the cell-intrinsic mechanisms underlying axonal reorganization after injury, without the need to sacrifice the animals. We focused on EPB-rich axons, known to be affected by aging^9^ to investigate whether their response to injury differs from that of the young adult brain. Using time –lapse imaging from 4 days pre-lesion to 3 months post-lesion, we examined regeneration, degeneration, and synaptic reorganization. Our findings revealed impaired regeneration in the aged brain, along with increased synaptic dynamics after injury. A computational model allowed us to explore whether these dynamics could be compensatory, showing that they may allow for partial recovery in a memory task following injury. This approach provides valuable insight into cellular responses to axonal damage during aging and establishes a foundation for investigating potential therapeutic targets in the aged brain.

## Results

To assess the responses of cortical axons and their synapses to local injury, we locally increased the power of the multiphoton imaging femtosecond laser^17^. The typical axonal response to injury and the imaging schedule are shown in Fig. 1. Typically, axons of appropriate length and type are selected pre-lesion (Fig. 1 A). During the lesion day (day 0), the axon of interest is lesioned by laser-mediated axotomy (Fig. 1 B, E). EPB-rich axons (L2/3/5 and thalamocortical axons) were then imaged up to 3 months post-lesion to study regeneration, degeneration and synaptic dynamics (Fig. 1 C-F). We followed EPB-rich axons (*n* = 14, all experiments) over multiple time-points (Fig. 1F) o ver da ys and mo nth s with in vivo two-photon imaging. The identification of the same axons was done using anatomical reference points (e.g., vessels and bifurcation points). The location of the lesion (*n* = 14) along the axon was chosen to yield disconnected axons of comparable lengths to previous experiments, ∼150 − 500 µm length cut (on average 314 ± 111 µm). The visible part of the axon had a total length of ∼400-1500 µm (on average 738 ± 317 µm). Following the injury, there was a typical response, including the retraction from the lesion site (Fig. 1B-E; length of 18 ± 4 µm, 20 min post-lesion), and an auto-fluorescent mark left by the laser damage (Fig. 1E; diameter of 9 ± 2 µm, 5 min post-lesion). To confirm that the axon was cut, we examined the axon several time-points post-lesion and observed that the distal portion of the axon degenerated (*n* = 14). In all cases, the proximal portion remained and was imaged at later time-points to examine its retraction, regeneration and synaptic dynamics over time.

**Figure 1.**
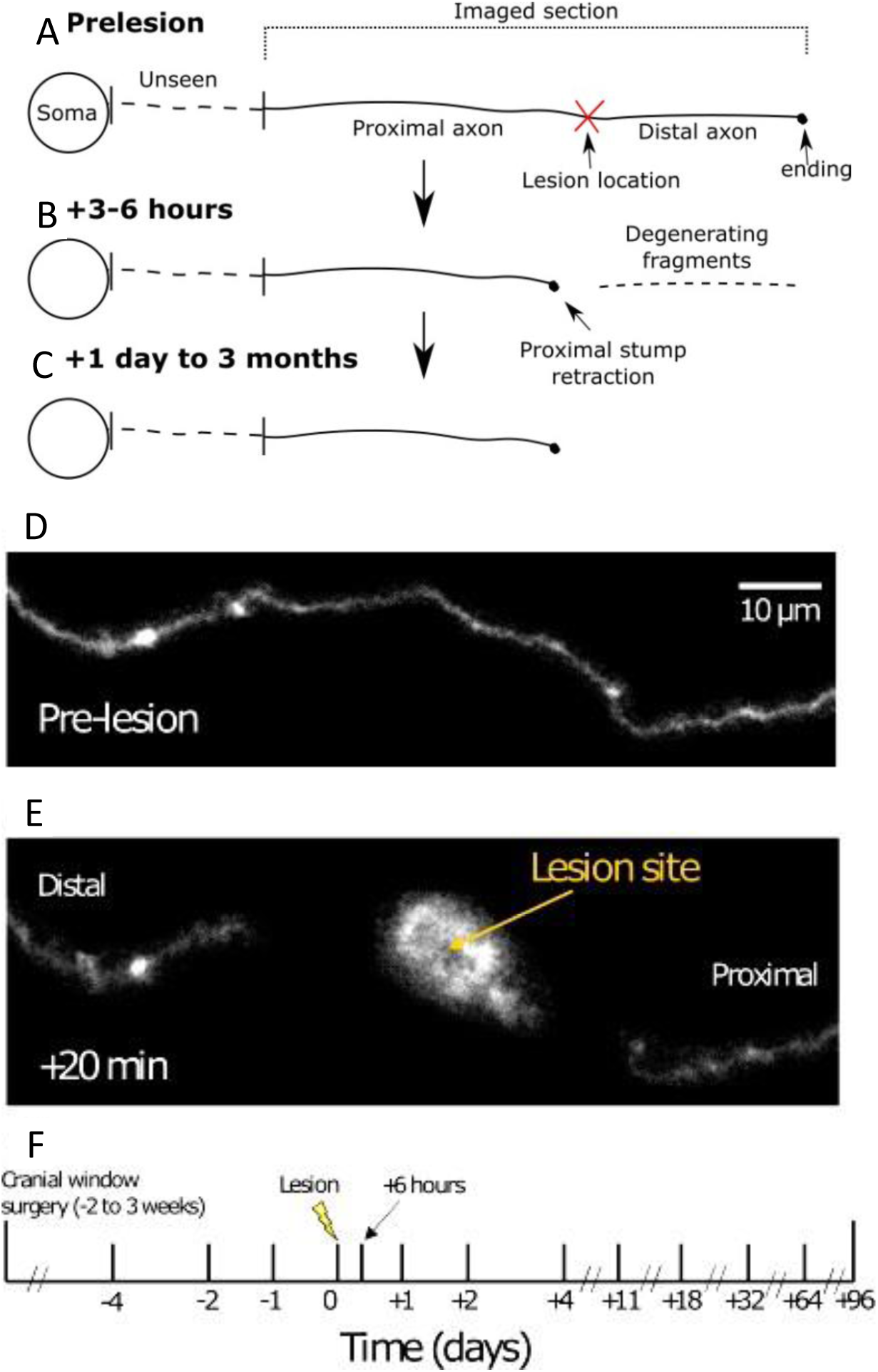
Tracking Axonal Dynamics Following Lesion in the Aged Brain: Degeneration, Retraction, and Regeneration. **A**, An axon of appropriate length is selected before the experiment begins. The axon is then measured, and lesion location is selected. The proximal portion of the axon (closer to the soma) is imaged for regeneration and retraction of axonal stump following the lesion. The distal portion is imaged until degeneration is complete. **B**, Typically by 6 hours post-lesion, fragmentation of the distal portion can be observed. **C**, Degeneration of the distal portion is usually complete by 1-day post-lesion. The proximal portion of the axon is imaged up to 3 months post-lesion for retraction, regeneration and synaptic dynamics. **D**, An example of an image taken before the lesion. **E**, A typical response captured at 20 minutes post-lesion. Both proximal and distal axons retract a comparable distance from the lesion site. **F**, Imaging schedule (days) to capture the degeneration, regeneration and synaptic dynamics following injury. Axons were imaged up to 3 months post-lesion.

### Axon Regeneration is not Observed after Injury in the Aged Cortex

The regeneration of EPB-rich axons was monitored in aged mice. No regeneration was observed from the proximal stump up to 3 months post-lesion (n = 9 out of 9 axons). However, in one case, a small growth was observed originating from a branch of an injured axon (16 µm over 3 days).

### Axon Retraction Rates in Aged Brain are Comparable to Non-regenerating Axons in young mice

Axons retracted 53 ± 26 µm on average up to 3 months post-lesion (*n* = 9; Fig. 2), of which most of the retraction occurred in the first 1 day (36 ± 25 µm), followed by slow retraction until 3 months post-lesion (6.8 ± 11 µm, *p* = 0.0078, Wilcoxon signed rank). Retraction dynamics in the aged brain (*n* = 9) was comparable to non regenerating axons in the young adult cortex (*n* = 53, *p* = 0.185, Kruskal-Wallis). Axons in aged vs young mice retracted (distance away from lesion site µm); at day 1, 36 *±* 25 vs 44 *±* 79 µm; at day 2, 38 *±* 25 vs 52 *±* 93 µm; at days 4-5, 39 *±* 25 vs 62 *±* 87 µm; and at days 8-11 43 *±* 25 vs 58 *±* 87 µm (aged and young adult^12,13^, respectively).

**Figure 2.**
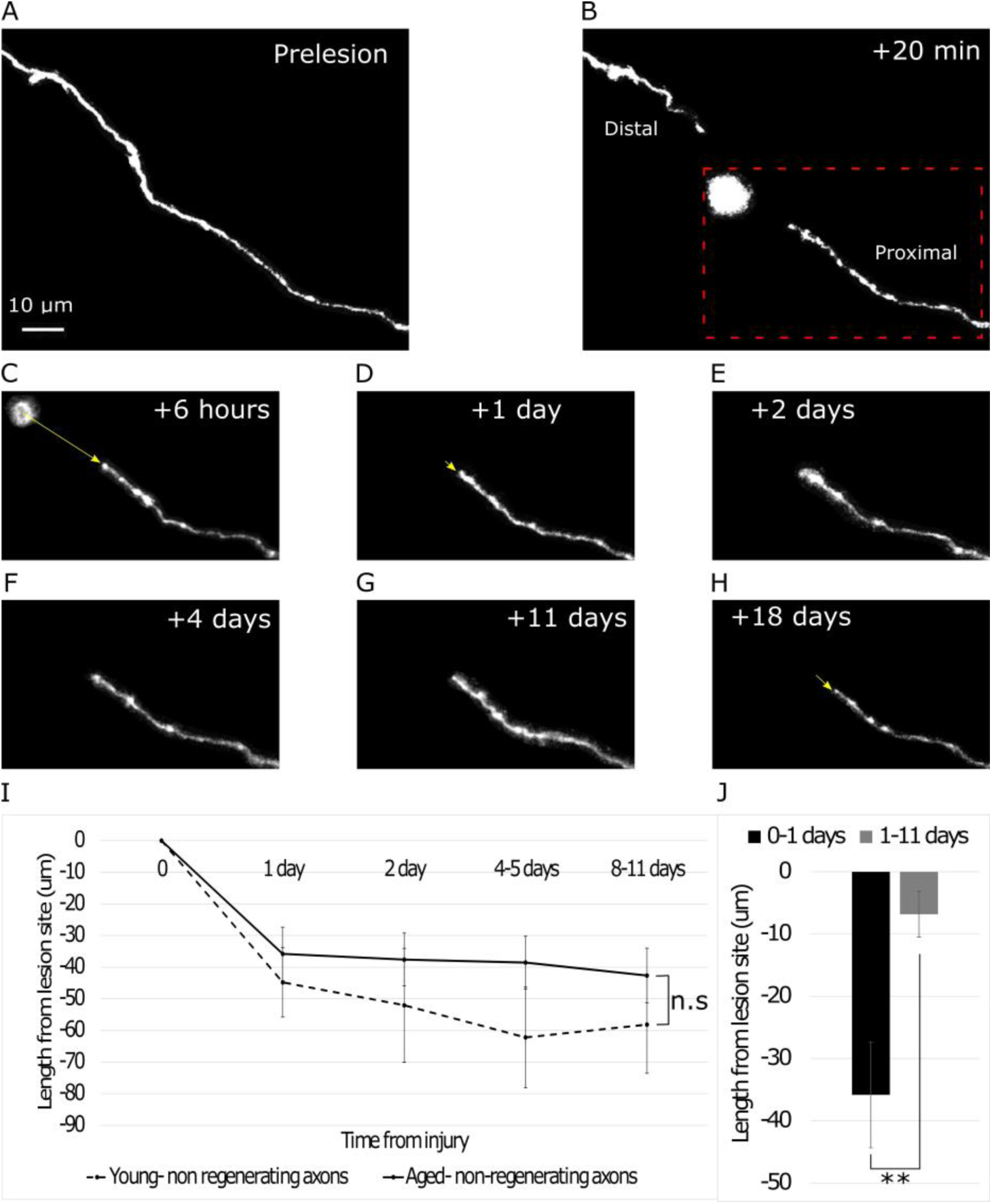
Axonal Retraction Dynamics in Aged vs. Young Animals Following Lesion. **A**, Two-photon image of a cortical axon in the aged brain pre-lesion. **B-H,** Images of the same axon 20 minutes, 6 hours, 1 day, 2 days, 4 days, 11 day, and 18 days post-lesion respectively. Yellow arrows indicate where the proximal stump retracted to. Red dotted box refers to the same location followed in C-H. **I**, A quantitative comparison of retraction in aged versus young animals (*p* = 0.185, Kruskal-Wallis). **J**, Retraction occurred in 2 phases; an acute phase which consisted of the most retraction (0-1 day post-lesion), and a chronic phase in which very little retraction occurred (*p* = 0.0078, Wilcoxon signed rank). *, *p <* 0.05; **, *p <* 0.01; Error bars, SEM.

### Axon Degeneration in Aged Brain is Comparable to the Young Brain

To capture the degeneration dynamics of EPB-rich axons in the aged mouse cortex, we imaged at several intervals post-lesion, until at least 50% of the distal portion of the axon degenerated. Acute Axonal Degeneration (AAD), a process of rapid (minutes) fragmentation occurring both proximally and distally from the lesion site, has been described in the spinal cord^18^, and optic nerve^19^, but was not observed in the cortex^12,13,20^. We did not observe AAD (*n* = 14), which is consistent with previous studies in the cortex of young adult mice.^12,13,20^ Around 85% (12 out of 14) axons degenerated completely by 1 day post-lesion, of which all degenerated at least 50% by 6 hours post-lesion (Fig. 2). The remaining 15% degenerated by day 2. For axons in which we captured fragmentation of the distal portion (WD; *n* = 6), fragmentation onset was between 1 hour to 3 hours post-lesion. Due to the lack of short interval time-points, it was not possible to do a quantitative comparison to the young adult counterpart.^12,13,20^ It appears from our qualitative comparison, however, that their dynamics are comparable.

### Increased Synaptic Dynamics Following Injury in the Aged Brain

To study synaptic changes over time in the aged, injured cortex, we imaged 8 axons (EPBn = 277) for 8 time-points before and after laser-mediated axotomy (–4 days, –2 days, –1 day, 0, +6 hours, +1 day, +2 days, +4 days). An example of typical bouton addition and elimination over time is shown in Fig. 4. We observed that the total number or density of EPBs significantly increases at +6 hours post-lesion (from 0.043 ± 0.012 at pre-lesion, to 0.048 ± 0.012 at +6h; n = 8, p = 0.0024, Friedman’s followed by Wilcoxon signed rank with Bonferroni correction; Fig. 5A; pre-lesion vs +6h, p = 0.0078, Wilcoxon signed rank with Bonferroni correction), and then it returns back to baseline density levels at +4 days post-lesion (0.043 ± 0.016; 6h vs 4d, p = 0.0078, Wilcoxon signed rank with Bonferroni correction). We next determined the TOR, comparing the pre-lesion (0.16 ± 0.028) time-intervals, and the [0 to 1d] (0.23 ± 0.07) time-interval, and found a significant increase (n = 8 axons, p = 0.039, Wilcoxon signed rank). Furthermore, when examining whether the increase in TOR was due to increase in gains or losses (Fig. 5C-D), we found no significant difference between gains and losses over one day, but there was a significant increase in total number of gains (0.29 ± 0.17) vs losses (0.1 ± 0.05) in the [0 to 6h] time-interval (p = 0.0028, Man-Whitney U; Fig. 5C). This indicates that there was an immediate reaction to injury, with a total increase in boutons (Fig. 5A-C) caused by a decrease in losses. Both gains and losses go back to baseline rates by 2 days post-lesion, i.e. there is no net gain or loss over time (Fig. 5A-C).

**Figure 3.**
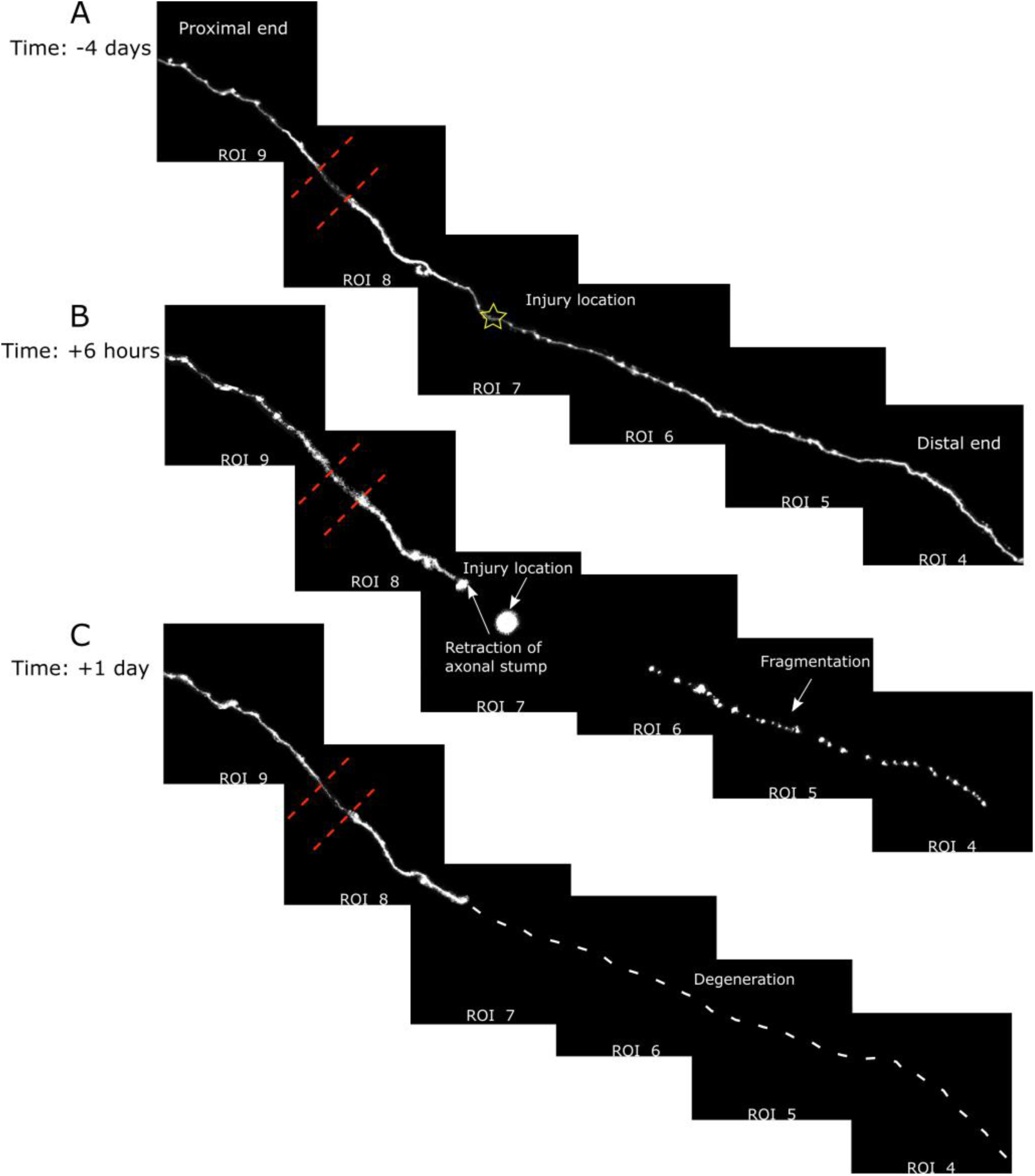
Example of axon degeneration in the aged somatosensory cortex. **A**, An example of an axon at a pre-lesion time-point, illustrating the location of the injury with a star. **B**, The same axon at 6 hours post-lesion, showing fragmentation in the distal part of the axon. **C**, At 1 day post-lesion, all of the distal part of the axon degenerated, shown by the white dotted line. The red dotted line indicates the location of a crossing vessel, causing reduced imaging intensity.

**Figure 4.**
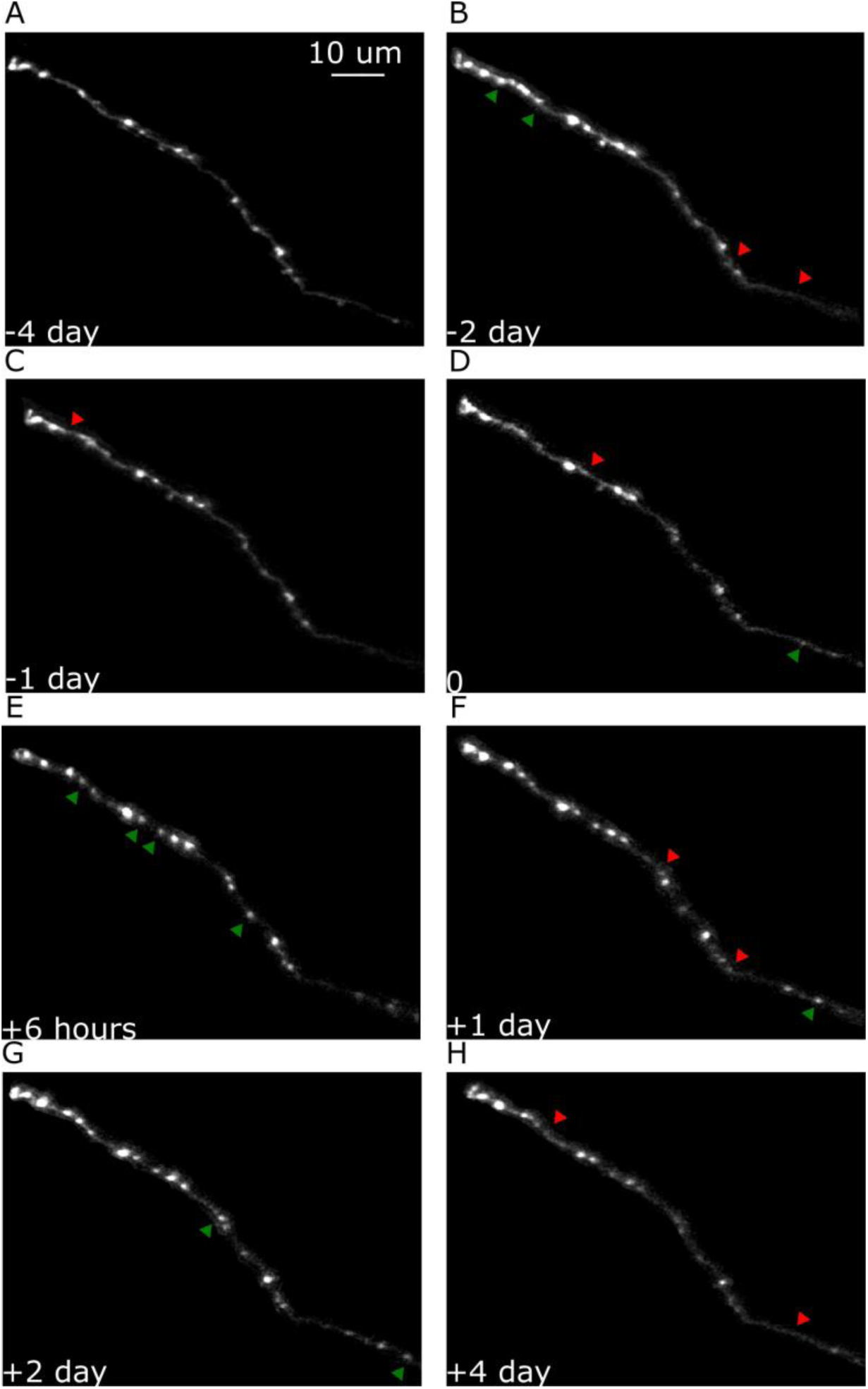
Two-photon *in vivo* imaging of EPB turnover in the aged mouse cortex. **A-D**, Pre-lesion EPB dynamics. **E-H**, Post-lesion EPB dynamics. **Green arrows**, added EPBs; **Red arrows**, eliminated EPBs.

**Figure 5.**
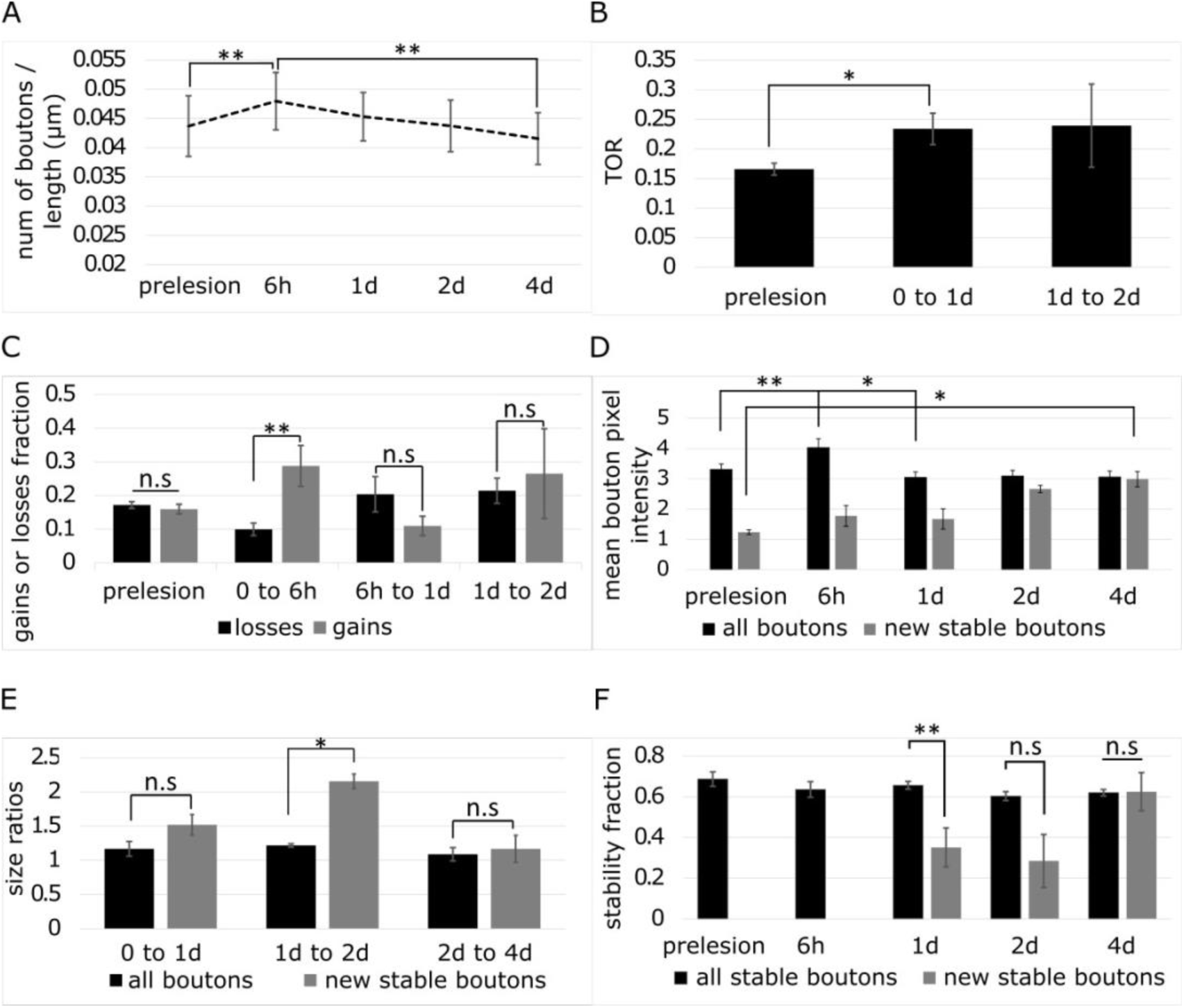
Quantification of EBP dynamics before and after injury in the aged somatosensory cortex. **A**, Density of boutons over time. **B**, The TOR of 1 day time-intervals. Prelesion is the average across the [-2d to –1d] and the [-1d to 0] time-intervals. **C**, Gains and losses fraction for different time-intervals. Prelesion is the average across all prelesion time-intervals, including [-4d to –2d], [-2d to –1d], and [-1d to 0]. **D**, Mean bouton pixel intensity across all time-points. Prelesion is the average across all prelesion time-points, including: –4d, –2d, –1d, and 0. **E**, The size ratios of boutons between 2 time-points. **F**, The stability fraction of all stable boutons, and new boutons that have stabilized following injury. Prelesion is the average across all prelesion time-points, including: –4d, –2d, –1d, and 0. TOR, turnover rate; *, *p <* 0.05; **, *p <* 0.01; Error bars, SEM.

We next studied the relative size changes of boutons pre– and post-lesion (Fig. 5D-E), a measure of synaptic strength.^9^ We plotted the normalised mean bouton intensity across all time-points, and the size ratios between two time-points. We found an increase at 6 hours, but it was not maintained over a longer time (bouton size at pre-lesion = 3.30 ± 0.52, 6h = 4.03 ± 0.81, at 2d = 3.04 ± 0.51, *p* = 0.0078, Wilcoxon signed rank with Bonferroni correction). However, new boutons that become stable increased their size until they reached average bouton size (new boutons size at pre-lesion = 1.23 ± 0.19, at 4d = 3.00 ± 0.67; p = 0.0156, Wilcoxon signed rank with Bonferroni correction), possibly indicating a re-wiring mechanism. Overall, there is also no difference in size ratios of all boutons when looking at 1-day intervals, suggesting that there is a homoeostatic mechanism which keeps the overall output of the neuron constant over a longer time-scale. We also studied the size ratios of new and stable boutons that appear after injury. Stable boutons are defined as those being present for at least two consecutive sessions. New boutons that are also stable increase more compared to all boutons in the [1d to 2d] time-interval (1.21 ± 0.27 all boutons vs 2.15 ± 0.26 new stable boutons at 1*/*2*d*; *p* = 0.014, Mann-Whitney U), and go back to baseline by day 4. When comparing the stability fraction of all boutons, and new boutons after injury, we find that they are comparable (0.62 ± 0.13 all boutons vs 0.62 ± 0.24 new boutons at 4*d*; *p* = 0.51, Mann-Whitney U; Fig. 6 F), suggesting that new boutons are just as likely to become stable as all other boutons.

**Figure 6.**
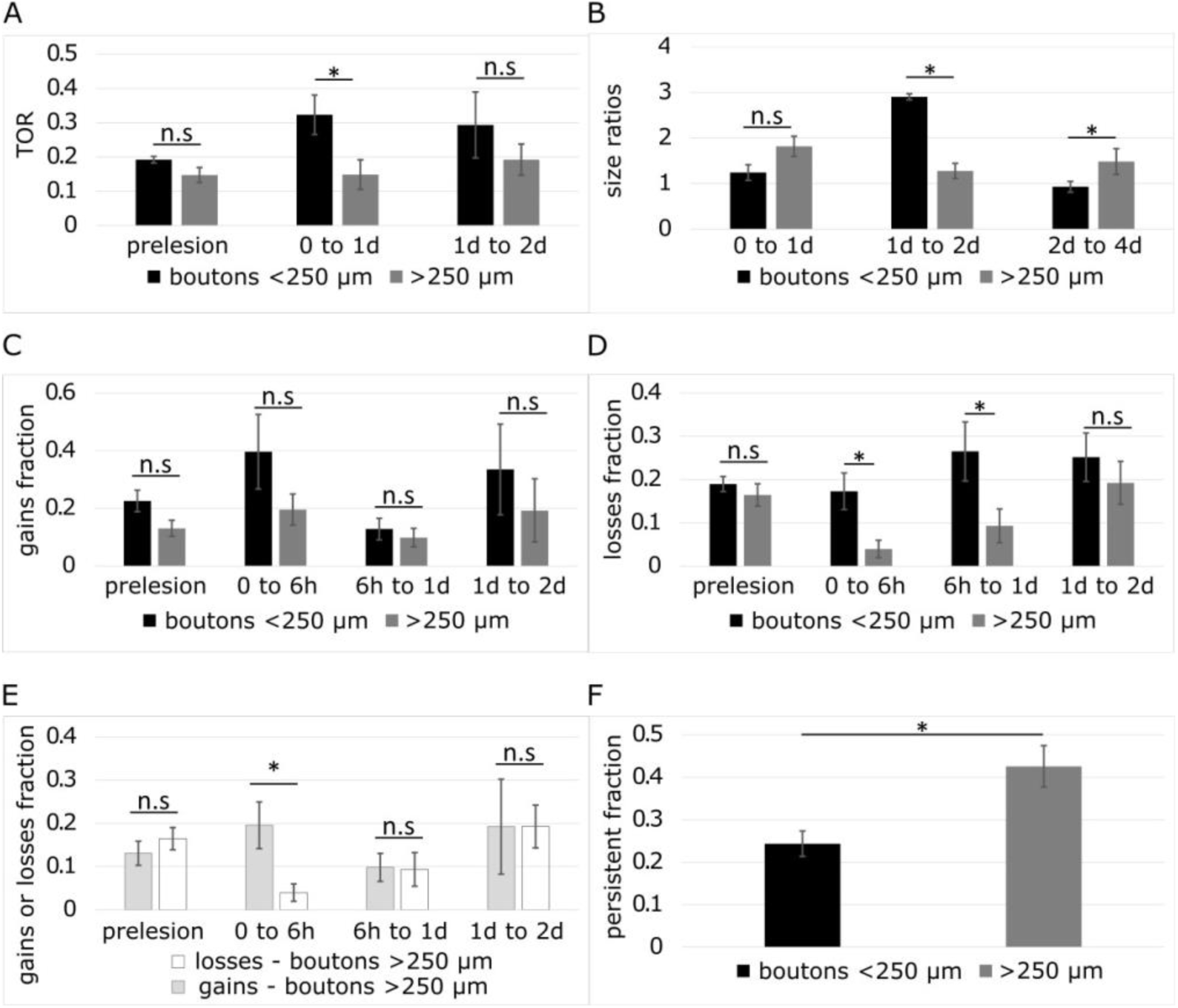
Quantification of EPBs close and far from the injury site. We split our EPBs into boutons close (*<* 250 µm) and far (*>* 250 µm) from the injury site. **A**, The TOR of 1 day time-intervals. Prelesion is the average across the [-2d to –1d] and the [-1d to 0] time-intervals. **B**, The size ratios of boutons between 2 time-points. **C-E**, Gains and losses fraction for different time-intervals, split into boutons close, and far from the injury site. Prelesion is the average across all prelesion time-intervals, including [-4d to –2d], [-2d to –1d], and [-1d to 0]. **F**, Persistent fraction for boutons close and far from the injury site. Persistent boutons were present in all the imaging session. TOR, turnover rate; *, *p <* 0.05; Error bars, SEM.

Overall, we found that after injury, there is an increase in synaptic dynamics in which boutons are gained and lost soon after injury, and these new boutons become stable and increase their size more than pre-existing boutons. This might indicate an attempt to compensate for the loss of synapses that were caused by the injury.

We next examined whether there are differences in the dynamics of boutons which are close to the injury site (Fig. 6), defined as *<* 250 µm (which we refer to as close boutons, or *EPB_close_*), and boutons far from the injury site, *>* 250 µm (which we refer to as far boutons, or *EPB_far_*). When looking at turnover rates of close vs far boutons (number of *EPB_close_* = 134, number of *EPB_far_* = 143), we observed that the portion of axon closer to the injury site is affected to a greater extent than the portion that is further away. This is indicated by an overall higher TOR at [0 to 1d] time-interval (0.32 *±* 0.16 *EPB_close_* vs 0.14 *±* 0.12 *EPB_far_*, *p* = 0.039, Mann-Whitney U), and that there are more non-persistent boutons (0.75 *±* 0.08 *EPB_close_* vs 0.57 *±* 0.13 *EPB_far_*, *p* = 0.019, Mann-Whitney U) and less persistent boutons (0.24 *±* 0.08 *EPB_close_* vs 0.42 *±* 0.13 *EPB_far_*; *p* = 0.013, Mann-Whitney U). Overall, there are higher losses in *EPB_close_* vs *EPB_far_* at [0 to 6h] (0.17 *±* 0.12 *EPB_close_* vs 0.04 *±* 0.06 *EPB_far_*; *p* = 0.04, Mann-Whitney U) and [6h to1d] (0.27 *±* 0.19 *EPB_close_* vs 0.09 0.1 *EPB_far_*; *p* = 0.038, Mann-Whitney U). In addition, there is no difference between fraction of gains vs losses in close boutons (*p* = 0.2, Mann-Whitney U), but there are more gains vs losses in boutons far from the injury site (0.2 ± 0.15 gains vs 0.04 ± 0.06 losses at [0 to 6h]; *p* = 0.04, Mann-Whitney U). Boutons close to the injury site (stable after injury) showed the greatest increase in size at [1d to 2d] (2.91 *±* 1.02 *EPB_close_* vs 1.27 *±* 0.45 *EPB_far_*, *p* = 0.031, Mann-Whitney U). And boutons far from the injury site (stable after injury) showed an increase in size that was marked at [2d to 4d] time-interval (0.93 *±* 0.15 *EPB_close_* vs 1.5 ± 0.33 *EPB_far_*, *p* = 0.031, Mann-Whitney U). This indicates that while boutons close to the injury site are more dynamic after injury, more of these boutons are lost, whereas less boutons far from the injury site are lost and are more stable over time. This might occur in response to injury as an attempt to re-wire or compensate for the loss of boutons caused by the injury. Overall, our data indicate that there might be a re-wiring mechanism after injury, in which more boutons close to the injury site are lost, and more boutons far from the injury site are gained, which results in no net gain or loss in boutons by day 4 post-lesion. Boutons close to the injury site increase their size the most (∼3X), and new boutons become stable over time after the injury.

### Computational Modelling of Injury in the Aged versus Young Brain Model outline

To investigate whether synaptic reorganization observed in the aged brain can elicit recovery after injury, we decided to use a computational model. Unlike recurrent neural networks in the machine learning literature, we use a recurrent neural network aimed at mimicking biological neurons, which are spiking, and learn information via a Hebbian learning rule, which is more biologically realistic than methods often used in machine learning, such as back-propagation. The model is composed of 100 integrate and fire excitatory neurons^21^ which are recurrently connected (Fig. 7B), with a Hebbian learning rule. We examine whether this network can recover from injury in a computational model, described below. Our aim was to study, 1) whether the changes in synaptic dynamics (TOR) or regeneration, change the capability for memory recall, 2) whether an injury would impair recall, 3) and whether there can be recovery following injury.

**Figure 7.**
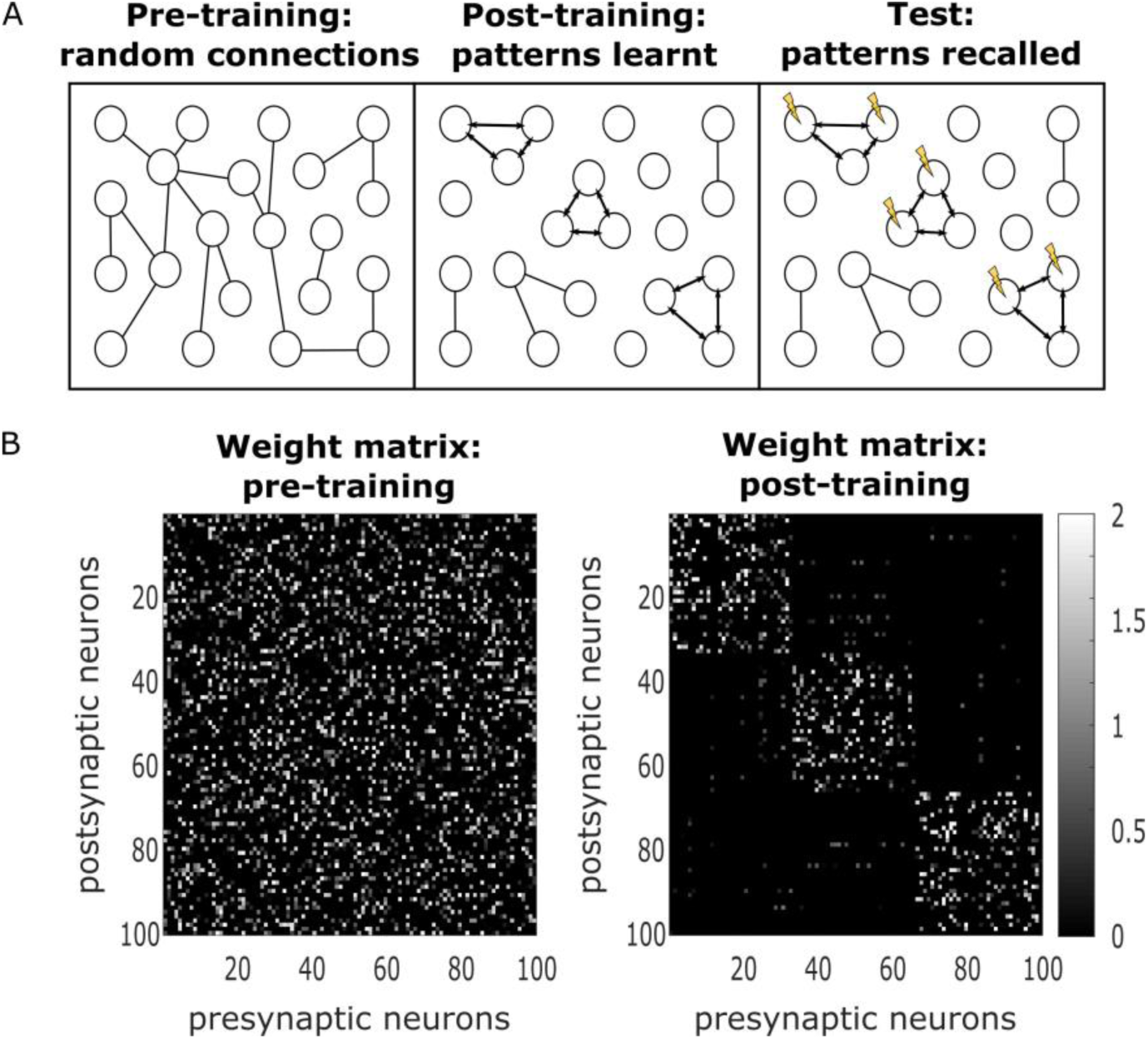
Modelling illustration and weight matrix evolution. **A**, Weights were initialized randomly according to connectivity observed in the somatosensory cortex (left). We trained “patterns” using external stimulation delivered to 3 sets of neurons (*N_pattern_* = 33). The result was 3 highly connected patterns (middle). We use these “patterns” in an associate memory test to observe how well the network has learned from the training, by stimulation ∼60% of the neurons in each pattern (right), and computing a score based on the activation of the neurons that were not stimulated (i.e. “test neurons”, *N_test_* = 13). **B**, Example of randomly generated weight matrix pre– and post-training. The white squares represent strength of connection between a pre– and post-synaptic neuron. Black represents no connectivity between the neurons.

This model was used to study whether changes in synaptic dynamics would affect memory storage. To model memory storage in the cortex, we trained the model to have “patterns”, which represent different encoded memories. These patterns are groups of highly connected neurons (Fig. 7B), which evolve during training by repeated stimulation of external current (an example of one training simulation is shown Fig. 8A). During training, we recorded *W*, a weight matrix containing the synaptic strength of connections between neurons, and found that neurons in the same pattern were strongly potentiated (Fig. 7B). Sparse noise in form of external current, is also presented during training (Fig. 8A, left). After training, to test how well those patterns or memories are stored, we use an associate memory task (Fig. 7A, right). During testing, we stimulate ∼60% of the neurons in each pattern (20 out of 33 of the neurons) with high external current (Fig. 8B, left). We then compute the recall performance of each pattern, by examining whether the neurons that were not directly stimulated in the test (13 of the 33 neurons), were activated by the test current (Fig. 8B, right). Finally, in the injury model, in which we tested how the young and aged brain respond to injury (using parameters from the injury experiments), we compared performance in aged EPB vs young EPB conditions, and aged EPB vs young TB conditions, to study whether recovery is possible.

**Figure 8.**
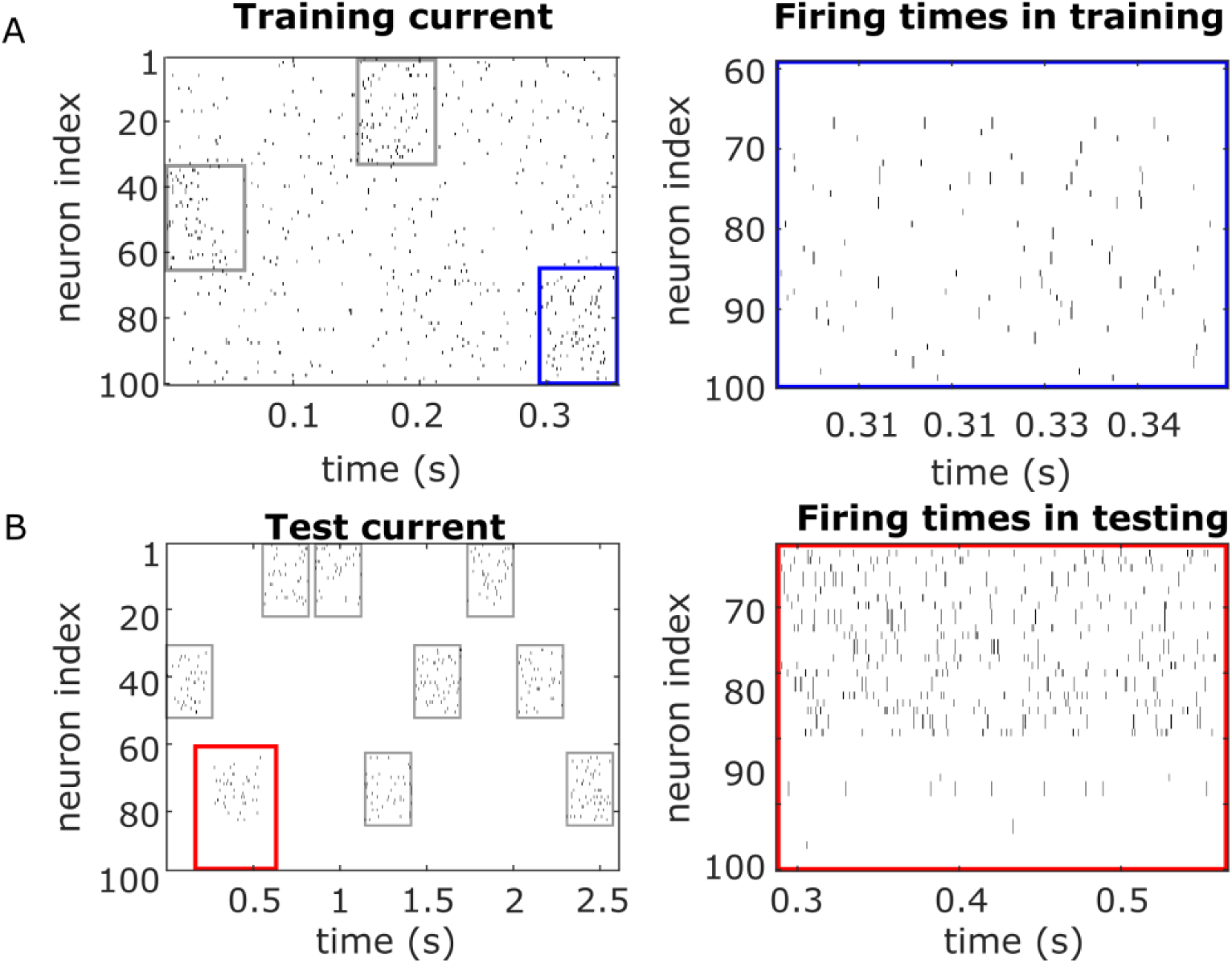
Training and testing in the computational model. We performed our modelling simulation using a recurrent neural network using the Integrate and Fire model with Hebbian plasticity. **A**, An example of 1/4 training simulation. We trained “patterns” using external stimulation delivered, *reps_train_* = 4 times (the figure shows only 1 rep), to 3 sets of neurons (*N_pattern_* = 33), for a total simulation length of *Sim_train_* = 1.8*s*. The result was 3 highly connected patterns. Average firing rates, 10 Hz and 26 Hz (EPBs vs TBs); Maximum firing rates, 80 Hz and 148 Hz (EPBs vs TBs). **B**, An example of an entire testing simulation (*Sim_test_*= 2.61*s*). Each pattern was tested *reps_test_* = 3 times. We observe how well the network has learned from the training, by stimulation 60% of the neurons in each pattern, and computing a score based on the activation of the neurons that were not stimulated (i.e. “test neurons”, *N_test_*=13). Average firing rates, 57 Hz and 50 Hz (EPBs vs TBs); Maximum firing rates, 184 Hz and 165 Hz (EPBs vs TBs). **Black lines**, neuron spikes or firing times; **Grey boxes**, stimulation times; **Blue box**, location for firing times in training; **Red box**, location for firing times in testing.

### Baseline Model Results

We observed a few interesting properties from our baseline simulations (i.e. training patterns without applying injury). We found comparable results in the analysis of both our EPB and TB simulations. We were first interested in how the learning rate *β* affects the turnover rate (TOR) in our model, and found that *β* is positively correlated with TOR (*n* = 50, *r* = 0.97, *p* = 3*e^−^*^30^, Pearson’s correlation), and error (*n* = 50, *r* = 0.81, *p* = 5*e^−^*^1^^3^, Pearson’s corr*−*elation). Higher rates of TOR, which are representative axons in the of aged brain, are also positively correlated with error (*n* = 50, *r* = 0.84, *p* = 2*e^−^*^1^^4^, Pearson’s correlation). This indicates that the performance in this memory task decreased with higher rates of TOR, which is comparable to what is found in the aged brain experimentally.^9^ This might occur because while higher learning rates lead to forming connections more quickly, they also lose (i.e. forget) connections more quickly, and thus are less stable over time. As noise is also presented between training stimulations, it is likely that higher learning rates lead to fast forgetting of the patterns. This is supported by the observation that higher TOR is inversely correlated with connectivity (i.e. lower weights) in the patterns (*n* = 50, *r* = 0.94, *p* = 2*e^−^*^2^^4^, Pearson’s correlation). We next investigated how the model of the young and aged brain performs in baseline conditions (i.e. without applying an injury, Fig. 9A-C). For the comparison, we fitted parameters for aged EPB, young EPB, and young TB conditions (see parameter fitting in methods for further details), and computed their performance in a memory task after training. All simulations were repeated 3 times, and the averages are reported here. We found that the young EPB condition performs the best when compared with aged EPBs (Fig. 9C; 0.37 ± 0.06 error – young EPBs; vs 0.52 ± 0.02 error – aged EPBs, *n* = 50*, repeats* = 3*, p* = 7*e^−^*^1^^4^, Mann-Whitney U). This is supported by results from prior work, which found that there is reduced performance in cognitive tasks in the aged brain.^9^ Interestingly, in our model TBs were found to have less optimal dynamics to store patterns in our memory task (Fig. 9A; 0.47 ± 0.09 error – young EPB, vs 0.77 ± 0.05 error – young TB, *n* = 50*, p* = 7*e^−^*^1^^8^, Mann-Whitney U). This is consistent with experimental studies in which EPBs are thought to encode for long-term memories, because they are bigger and longer lasting, whereas TBs are thought to have a more supporting role, as they are smaller and more dynamic (see^22^ for a review of the topic).

**Figure 9.**
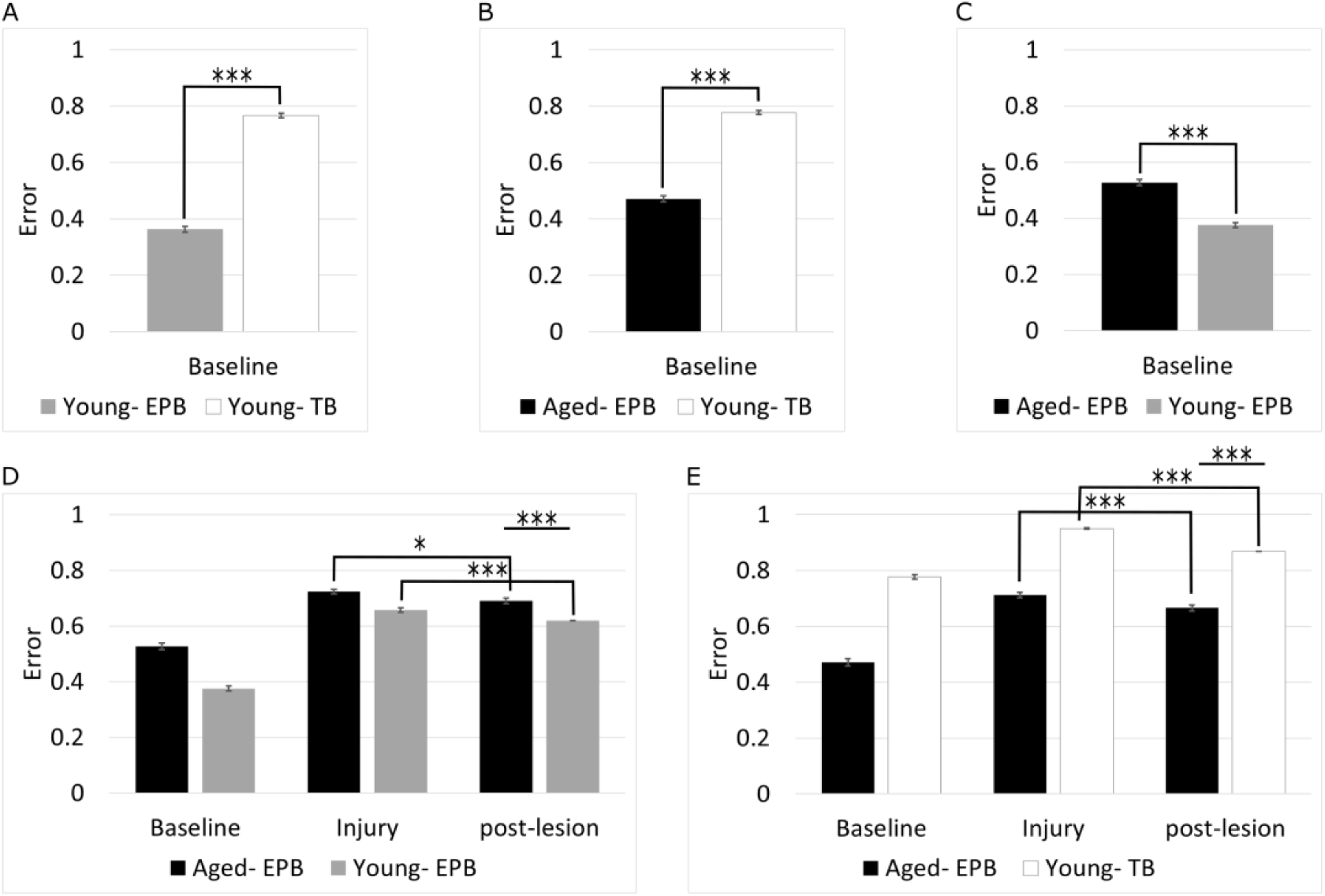
Modelling results. We compared aged EPBs (*n* = 50), young EPBs (*n* = 50), and young TBs (*n* = 50) to determine which would perform better in baseline and following injury. **A-C**, EPBs are significantly better than TBs in performing a memory task (A-B). In addition, young EPBs have more optimal dynamics for storing memory, than aged EPBs (C). **D-E**, We compared the different conditions following injury, and found in all cases a large and significant increase in error after the injury. In general, we found a small recovery from injury in all conditions, with young EPBs having the best performance post-lesion (0.62 0.06 vs 0.69 0.08, young EPB vs aged EPB, *p* = 2*e^−^*^6^, Mann-Whitney U). We also found that young TBs recover the most from injury (0.08 vs 0.04 recovery, young TBs vs aged EPBs), but still perform worse overall (0.86 0.03 vs 0.66 0.08, young TBs vs aged EPBs, *p* = 1*e^−^*^1^^7^, Mann-Whitney U). Error bars, SEM.

### Injury Model Results

We next examined how our model of the aged and young brain respond to an injury, as is done experimentally (for details for injury model see Methods, Fig. 10). As expected, we found that there was an increase in error following injury in all conditions (Fig. 9D-E). We found that young EPBs have better capacity for memory storage than aged EPBs, and that in general EPBs have more ideal dynamics than TBs in storing memories.

**Figure 10.**
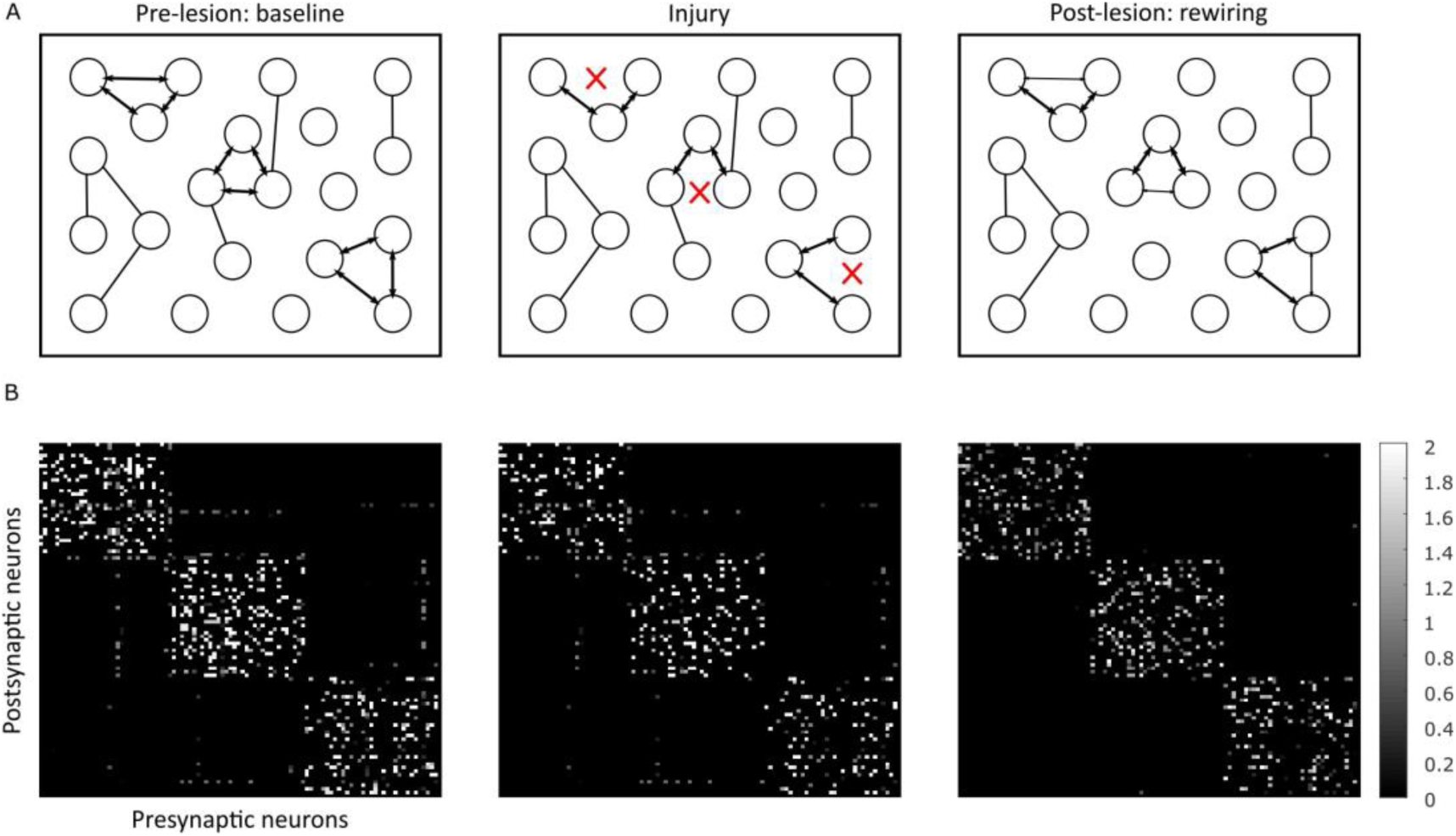
Injury model. **A**, An illustration of the experiment performed to test parameters from the ageing and injury results showed in this paper. Left-as in training, we trained a network using parameters from the somatosensory mouse cortex, and fitting TOR values from results in this paper and prior work.^9,12,13^ Middle-to model the injury we performed experimentally, we cut a number of connections, consistent with results from this paper. Right-we then add and remove connections, in a manner that reflects the experimental observations of the aged cortex. We perform a test to compare memory performance at each of these stages. **B**, An example of weight matrix generated at each step for the EPB aged condition; starting with baseline on the right, to injury, and post-lesion to the left. After the weight matrix is generated, 1 repeat of training stimulation is applied, followed by testing stimulation. TOR, turnover rate.

Following injury, we observed a small but significant recovery in the aged EPB model in all simulations (Fig. 9D-E). When comparing to young EPBs, we found that aged EPBs perform worse after injury (Fig. 9D; 0.69 ± 0.08 in aged EPBs, vs 0.62 ± 0.06 in young EPBs, *p* = 2*e^−^*^6^, Mann-Whitney U). The young EPBs have overall the best performance post-lesion compared to all other conditions. This suggests that the young EPB dynamics are better at recovering from injury than aged EPB dynamics. In the model of the young TB condition (Fig. 9E), the highest recovery from injury was found (0.86 ± 0.03 vs 0.66 ± 0.08, young TBs vs aged EPBs, *p* = 1*e^−^*^1^^7^, Mann-Whitney U), with 0.08 reduction in error, which is closer to its baseline performance than all other conditions (0.09 worse than baseline, compared to aged EPB which is 0.19 worse than baseline). However, the error is still significantly higher than both young and aged EPBs, demonstrating that even with better regeneration, TBs are still worse at storing memory than EPBs. To summarise, in our modelling simulations, we investigated how the young and aged brains respond to injury, and revealed that while the young brain can recover the most by regeneration, the aged brain can also attempt a small but significant recovery by synaptic re-wiring.

## Discussion

In this study, we used *in-vivo* two-photon imaging through a cranial window on the somatosensory cortex to examine the response of axons to localised microlesions for a period of up to 3 months post-lesion. As laser mediated axotomy induced small localized lesions, and are known to cause minimal glial scar in the cortex, it is unlikely that these lesions induce a strong immune response, or prominent scarring.^12,13^ We followed the axons for at least two days post-lesion, and were able to confirm in all cases that the correct axons were injured. The aim of this study was to determine whether there are differences between the response of aged and young adult cortical axons to laser mediated microlesions. Because it has been shown previously that L2/3/5 and thalamocortical (TCA) EPB-rich axons are most affected by ageing, we restricted this study to these axons. Indeed EPB-rich axons display increased synaptic dynamics immediately following injury, which later return back to baseline levels. This response might indicate a re-wiring mechanism in response to injury. We also investigated the degeneration, retraction, and regeneration following a lesion *in vivo*. We found that degeneration of the distal portion was completed by the 1st day post-lesion in 85% of the cases, of which 71% of axons degenerated at least 80% (Fig. 3). Qualitatively, the degeneration dynamics we observed were similar to axon degeneration in the young-adult cortex.^20^ Quantitative analysis was not possible in this case, as there were not enough comparable time-points. Retraction of the proximal stump was also studied, and was found to occur in two phases (Fig. 2); an acute phase occurring in the first day, and a chronic phase continuing up to the last observed time-point (up to 3 months). Interestingly, these dynamics are similar to retraction rates reported for non-growing axons in the young adult cortex^12,13^ (*n* = 9 aged, *n* = 53 young).

Finally, we did not observe any regeneration in the aged brain, compared to 20% in young adult EPB-rich axons.^12,13^ In previous experiments in young adult mice (2-3 months old)^12,13^ it was observed that 20% of EPB-rich axons (from L2/3/5 and thalamus) regenerated over days and weeks. It is possible other cell types may show higher regeneration capacity in the aged brain, as shown in young adults^12^. Indeed, impaired regeneration of axons in the aged brain was expected, and is consistent with th e fact that the aged brain has significantly reduced ability to recover from disease and injury^6,7,23,24^ There are several factors that could affect the regeneration rate of cortical axons in the brain, and these are split into intrinsic and extrinsic factors.

### Extrinsic and Intrinsic Mechanisms for Impaired Regeneration in the Aged Brain

For extrinsic factors, the first possible culprit of impaired regeneration, is that the remaining distal portion of the axon impedes growth by blocking the path for regeneration. For instance, slow degeneration of the distal portion of the axon following injury, and the clearance of axon fragments has been investigated in the PNS and spinal cord.^12,13,24–27^ Evidence from PNS suggested that slower debris clearance of degenerated axons leads to impaired regeneration.^24,27^ Also, regeneration is improved when degeneration (of the distal axon) is induced before or simultaneously with an axon lesion.^25,26^ However, slowed degeneration is unlikely to be the cause of reduced regeneration in this case, as the axon degeneration in the aged cortex (Fig. 3) is comparable to young adult mice.^12,13,20^

Another extrinsic component that might influence regeneration is the presence of astrocytes, microglia cells, and glial scar. Currently, there is some controversy about whether these impede or aid regeneration. For instance, while astrocytic scar was long thought to have a detrimental effect on axon regrowth, a recent study provides direct evidence for improved axon regeneration associated with astrocytic scar.^28^ This is also supported by a study in the cortex, demonstrating serotonergic axonal growth in the presence of a glial scar environment.^29^ However, since laser-mediated microlesions have consistently reported no significant glial scar formation^12,13^, this is unlikely to have affected regeneration in this case. Microglia cells could also be a contributing factor for axon regeneration. Several different studies reported that microglia have age-related alterations^30^ and these alterations might cause a dysregulated response to injury.^31^ Their role in injury might be to clear debris remaining from the distal portion of the axon.^27^ Another possibility for impaired regeneration is based on intrinsic mechanisms. While we do not have any direct evidence indicating an intrinsic mechanism, we observed that the retraction rate of the proximal stump was comparable to non-regenerating axons (Fig. 2). Increased rate and magnitude of retraction has been inversely correlated with growth in juvenile and young adult mice^12,13^. A similar response was shown in another study, in which serotonergic neurons that lacked retraction had enhanced regrowth following injury, even in the presence of glial scar.^29^ This indicates that if an axon retracts less, it is more likely to regenerate, which is most likely related to the intrinsic response of axons to injury. It is possible that intrinsic changes in the biology of the axons that occur with ageing could also contribute to impaired regeneration. For example, growth cone generation has been suggested to be important for axon regeneration.^32^ In the cortex, a full growth cone is usually not present following injury, but instead we observe a small bulbous axonal stump^12,13^ (Fig. 2). Molecular regulators of axon growth could also affect regeneration capacity following injury, such as GAP-43^33^, PTEN and others.^34^ It will therefore be important, in future work, to identify which intrinsic factors might cause the reduced regeneration in aged mice, as they could be potential targets for therapeutic intervention, or for improving regeneration.

### Synaptic Dynamics Following Injury in the Aged Brain – a Maladaptive or Compensatory Mechanism?

It is unknown why re-wiring might happen after injury in the aged brain, but it is possible that it is an attempt to compensate for boutons lost on the distal part of the axon (unlike in the young brain in which axon regeneration is observed, Fig. 1). There are two competing theories on why synaptic dynamics might change after injury; either the increased synaptic dynamics are a compensatory mechanism, or are a maladaptive response. It is possible that increasing synaptic dynamics could be a maladaptive response to injury, or a dysfunctional state. For instance, increasing levels of TOR are consistently reported in both boutons and spines (with spines having higher TOR than boutons^35^) with cognitive impairment^9,36^, injury^12,13^, ageing^9,10^, and with sensory deprivation.^37^ A study^38^ examining bouton dynamics following the induction of Long-Term Depression (LTD) found a decrease in bouton size, and an increase in bouton TOR, indicating that decrease in connectivity is associated with increased bouton TOR. Indeed, we also find in our baseline modelling simulations that higher levels of TOR are associated with decreased performance in a memory task, both in EPB and TB-like axons. However, since the increased synaptic dynamics in response to injury were short lived, and returned to baseline levels by day 4 post-lesion, it is unlikely that this change is maladaptive in the long term. Alternatively, it is possible that increased synaptic dynamics might be a mechanism of ensuring efficient learning even in old age^39^ or to compensate for the lack of regeneration observed in aged mice in this study. This theory is supported by several pieces of evidence. In our data, for instance, the net gain of boutons and size increase at 6 hours post-lesion (Fig. 5) might be an attempt to compensate for the boutons lost on the distal side of the axon, by forming new connections with neighboring neurons, or reinforcing existing connections. While there is a loss of boutons at 4 days post-lesion (with total bouton densities returning to baseline levels), many (60%, Fig. 5F) of the newly formed boutons are kept. Indeed, many of the new boutons increase their size more than all other boutons post-lesion (Fig. 5E), and also become stable (Fig. 5F). This might indicate a pruning mechanism, in which the network attempts to compensate for lost boutons and/ or impaired regeneration by optimising the available resources. To compensate for this, aged axons might rely on increased synaptic plasticity to cope with injury. Further evidence for re-wiring was shown by separating boutons into those close and far from the injury site (Fig. 6). Boutons close to the injury site were affected the most by the injury, having higher rates of TOR (with higher losses), and lower persistence. From the quantitative analysis, we observed that while more boutons that are close to the injury sites are lost, more boutons far from the injury site are gained at 6h post-lesion. It is possible that this is due to a pruning mechanism in which boutons that are less necessary after injury are lost (or alternatively are removed because of the inhibitory environment, or microglia^40^), and new boutons are formed to compensate for that loss, for continued effective coding. As revealed by a recent study, post-lesion re-wiring might occur due to enhanced excitability of pre-synaptic neurons onto the injured neuron^41^, which might aid recovery and maintenance of homoeostasis following an injury. In addition, it is possible that other structural or functional changes in nearby neurons occur to aid recovery, which we cannot observe with the current techniques, since we only imaged the injured axon, and we have not measured the electrophysiological properties of the injured axon or nearby neurons. It is possible that since we only make a small, minor lesion, that significant re-wiring or regeneration is not necessary for complete recovery from injury, as neurons are highly connected in the cortex. Alternatively, complete recovery via axon regeneration or synaptic re-organisation might be unachievable in the cortex. Imaging both pre– and post-synaptic dynamics after injury would be an elegant way to confirm whether there are other mechanisms at play.^42^

### Partial Recovery Following Injury in a Computational Model

Finally, we investigated whether the increased synaptic dynamics observed in this study can aid recovery from injury, using a computational model of recurrent excitatory integrate and fire neurons with Hebbian plasticity. We used intrinsic electrophysiogical parameters from the somatosensory cortex, and had the synaptic dynamics (TOR) fitted to the aged (this study, and^9^) and the young brain.^9,12,13^ We compared a model containing parameters of EPB-rich axons from the aged brain, to models based on a young brain with two different dynamics, using an associative memory task (in which neurons were trained before the injury is induced). One model was based on EBP-rich axons from the young brain, which regenerated only 20% of the time. The second was based on TB-rich axons, which have higher baseline TOR, and regenerate considerably more (55%). We found that in the aged brain, the synaptic re-wiring following injury, allowed a small partial recovery (Fig. 9D-E). Young EBP-rich axons recovered more after injury, and had the best performance compared to other conditions (Fig. 9D). Young TB-rich axons, recovered by the largest amount from injury compared to all other conditions, but still were significantly worse in the memory task than both young and aged EPBs following injury (Fig. 9E). This suggests, that according to our model, younger brains have better memory recall, (Fig. 9C, also supported by experimental findings^9^) the young brain can partially recover by regenerating the injured axon, and the aged brain can partially recover from injury using synaptic re-organisation (Fig. 9D-E). One interpretation of these findings might be that the young and aged brains have different mechanisms to cope with injury, one by regenerating injured axons and the other by re-wiring synapses to compensate for lack of regeneration. TB-rich axons in the young brain had the largest recovery, but still had worse performance in the memory task (Fig. 9E), possibly because TBs have a more modulatory role, rather than be optimised to memory storage.^22^ It might be that the role of the regeneration of TB-rich axons following injury is to modulate other neurons, and thus would still have worse memory capacity than their EPB counterparts. In addition, since the recovery found was not enough to return to baseline levels of performance, it is possible that there are mechanisms other than the regeneration or synaptic re-organisation of the injured axon, that can aid recovery from injury. For instance, since we do not image the neurons which are connected to the axon we injure, we do not know whether they are also affected by injury. It is possible that there are other functional and structural changes that might aid recovery in the nearby neurons. Since we use a simple model, and do not consider all the factors involved in the aftermath of injury in the cortex, we do not know how these dynamics might affect the whole brain. This is beyond the scope of this study, but will be interesting to study these dynamics in future work, in a more complex model, that would be more representative of biological neurons, and injury.

Overall, in this paper we found that 1) the aged brain has impaired regeneration which is not caused by slowed degeneration. 2) There is synaptic re-organisation following injury in which boutons are gained and increase size, and later return to baseline levels. 3) Our computational models suggest that synaptic re-organisation following injury in the aged brain could be a compensatory mechanism to injury, and the young brain could recover via regenerating the injured axons. According to our model, both mechanisms only allow partial recovery from injury, and it is possible that full recovery from injury is unachievable in the cortex.

## Methods

### Experimental Methodology Animals

For all the experiments, we used aged male mice (Thy-1-GFP-M; *n* = 12, *>* 2 years old with cytosolic GFP expression^9,13^) on C57BL/6 genetic background. As described previously^13^ animals were housed with littermates, in standard, individually ventilated caging and maintained in a 12 hour light/dark cycle with free access to food and water. All experiments involving live animals were conducted by researchers holding a UK personal license and conducted in accordance with the Animals (Scientific Procedures) Act 1986 (UK) and associated guidelines.

### Surgery

Surgery was performed as described previously^9,13^. For all imaging experiments, a cranial window was implanted overlying the somatosensory cortex under anaesthesia (0.083 mg/g ketamine, 0.0078 mg/g xylazine). Intramuscular dexamethasone (0.02 mL at 4 mg/ml) was administered to reduce inflammatory response, and subcutaneous bupivacaine (1 mg/kg), a local anaesthetic. The skull was exposed using a dental drill, and a few drops of lidocaine (1% solution) were applied on its surface. The glass coverslip that seals the window was placed directly over the dura and the bone edges, with a thin layer of agarose in between, and sealed with dental cement. Mice were given 2-3 weeks to recover and for the cranial window to clear, before the start of the imaging. Post-operative care was done by personal license holders, and trained staff at the animal house. Animals were monitored continuously for the first hour after surgery, daily until fully recovered (minimum 72 hr) and then at least three times a week after that to ensure no adverse effects are seen. Post operation analgesia was given (e.g. buprenorphine) within 72 hours post surgery, as advised by the Named Veterinary Surgeon (NVS). A number of humane end points were observed, and mice were sacrificed by a schedule 1 method or non schedule 1 method, if two or more of the following clinical signs were present: piloerection, hunched posture, reduced activity, increased docility or aggression, weight loss up to 20% of body weight, dehydration persisting for 24 hours during fluid replacement therapy, altered respiration, or self-mutilation.

### Imaging

Imaging was performed similarly to^13^ with a few minor alterations, described below. A two-photon microscope equipped with a tunable Coherent Ti:Sapphire laser and Prarie software for image acquisition was used for all imaging experiments. Mice were anesthetized with Isoflurane, an inhalation anaesthetic, and secured to a fixed support under the microscope. To prevent dehydration in the eyes, Lacri-lube (Allergan) was applied. To regulate body temperature (37C) an underlying heat pad was used and rehydration administered with isotonic saline solution (i.p.) required during long imaging sessions. Depth of anesthesia was closely monitored by a video camera and regularly checking reflexes (toe pinch) and respiratory rates. An Olympus 4 X lens with a 0.13 numerical aperture (NA) was used to identify characteristic blood vessel patterns, to relocate axons used in previous imaging sessions. An Olympus 40x with a 0.80 NA water-immersion objective was used to acquire the images. For optical zoom 1, images were typically 300×300 µm field of view, 512×512 pixels, 0.588 µm per pixel for *x, y* planes, and 3 µm per pixel for the *z* plane. For optical zoom 4, images were typically 75×75 µm field of view, 512×512 pixels, 0.147 µm per pixel for *x, y* planes, and 1 µm per pixel for the *z* plane. A point spread function was characterised, for images with optical zoom 4, by a Full Width at Half Maximum (FWHM) values of 0.45×0.45, 2.5 µm (*x, y, z*)). Either a water repellent pen or Vaseline (pure Petroleum Jelly) was applied around the cranial window to stabilize the meniscus for the 40x objective. A pulsed 910 nm laser beam was used, never exceeding 70 mW on the back focal plane. Each imaging session typically lasted for 60-90 min, during which time up to 40 image stacks (1 µm step size for images that were analysed) were collected. Typically, 12 non-overlapping axons were selected for lesioning in each animal. For synaptic analysis *in-vivo*, we first identified long axonal stretches and then collected high-magnification images of the synapses along the axon shaft at 6h, 1d, 2d, 4d, weekly, and monthly intervals (Fig. 1 C).

### Laser mediated micro-lesions

We used a Chameleon Ti:Sapphire, using a 9 µm diameter binary mask, ∼1000 mW at the back focal plane, 800 nm, and 3.2 µs dwell time to induce localised injuries in cortical layer 1. A total of 13 axon branches were lesioned across all imaging experiments and were observed up to 3 months post-lesion (*n* = 6 mice, *>* 2 years old). The amount of distal axon removed was between ∼150−500 µm (on average 314 ± 111 µm; in case multiple endings were present, we considered the total length of the disconnected portion), leaving a minimum of 250 µm of the surviving axon for post-lesion analysis, as in previous experiments^13^. Identified axons were severed by inducing a lesion using a single circular spot scan. Lesions were induced in the upper layers of the cortex, to a depth of 50 µm beneath the pial surface. Lesions were made as far apart as possible (minimum 300 µm). Cortical axons were imaged at variable intervals (Fig.1C) before and after lesion for as long as window clarity was preserved, or up to 3 months. During one of the imaging sessions, overlapping low-magnification overviews of the labelled blood vessels were collected to help identification of⍰axons in the fixed brain.

## Data analysis

The two-photon images were processed using a custom made, semi-automated Matlab MathWorks package called EPBscore^43^(synaptic remodelling), and ImageJ (length quantification, file conversions). For further bouton dynamic analysis, we used a Matlab script to compute all discussed metrics. For bouton counting, we also used the bouton detection software from.^44^ Figures were prepared using Matlab, Microsoft Office Suite, ImageJ, Gimp and Inkscape. Axons were classified by morphological criteria, as previously described.^45^ Only EPB-rich axons were monitored in the imaging experiments.

### **-** Axon Degeneration

We calculated degeneration dynamics by first measuring the total length of the axon (proximal and distal), and then measuring the fraction of remaining length of the distal portion at the available time-points.

### **-** Axonal retraction

To calculate axon retraction, we measured the distance from the proximal tip to the lesion site at several intervals post-lesion. Growth was not visible in any axons that were imaged. We obtained measurements for 6 of the 7 axons that were lesioned, at days 0, 1, 2, 4, 11 and 32 post-lesion.

### **-** Axon Regeneration

To calculate regeneration, we measured the distance grown from the proximal stump between two time-points. We defined attempted regrowth if the elongation exceeded twice the maximum noise measurement (that is, > 6 µm; the maximum difference in measurements between fiducial points over repeated imaging sessions is 3 µm).

### **-** Axonal bouton quantification

EPB-rich axons were imaged and analysed. EPB rich axons have cell bodies that are located in either of L2-3, 5, or the thalamus, mainly form axo-spinous synapses and are relatively stable in the adult brain.^45^ The bouton metrics reported in this paper were calculated in the following way:

- EBPs close to the injury were selected based on the distance to the injury site. Close EBPs were *<* 250 µm away from the injury site, and far EBPs were *>* 250 µm away.
- Density of boutons over time was defined as: *Density*(*a*) = *n_a_/length_a_* (µm), where *n_a_* is the total number of boutons in session *a*. Pre-lesion refers to the average across the following time-points: –4d, –2d, –1d, and 0.
- Turnover rate (TOR) between two imaging sessions a and b, is defined as (*nG* + *nL*)*/*(2*N*), where *nG* and *nL* are the numbers of bouton gains and losses respectively, and *N* is the total number of boutons in session a. The daily TOR pre-lesion are calculated by averaging across [-2 to –1d] and [-1 to 0d] time-intervals.
- Gains and losses fractions were defined as: *Gains*(*a, b*) = *nG_b−a_/N* and *Losses*(*a, b*) = *nL_b−a_/N*.
- The size ratio (Δ*S*) of boutons between two imaging sessions a and b is defined as: Δ*S* = (*S_b_*)*/*(*S_a_*)), where *S_a_* is the size of a bouton at session a, and *S_b_* is the size of a bouton at session b. A ratio higher than 1 demonstrates an increase in size, whereas a ratio smaller than 1 demonstrates a decrease in size. We plot the average size ratios between different time-points before and after injury. Pre-lesion refers to the average across the following time-points: –4d, –2d, –1d, and 0.
- For the stability metrics, stable boutons were present for at least 2 consecutive sessions at a certain session. New boutons are boutons that first appeared after injury. New and stable boutons first appeared after injury, and were present for at least 2 consecutive sessions. Persistent boutons were present in all of the imaging sessions. Destabilized boutons were present for the first three sessions (for at least 2 sessions), and subsequently were lost before the last 2 imaging sessions (i.e. they were not present at all at the last 2 sessions).

## Statistics

We used SPSS or Matlab packages for all statistical analysis. For all statistical quantities in this paper, we first tested whether the data was normally distributed, and appropriate tests were used accordingly. Significance level was set for p*<* 0.05, and all tests were two-tailed. For multiple in comparison tests, we used a Bonferroni correction, where the significance level was equal to 0.05*/n*, where *n* was the number of tests performed (i.e. only p values under this are considered significant). For all figures, the error bars plotted are the Standard Error of the Mean (SEM).

## Computational modelling

All computer modelling was done in Matlab. We used a simple recurrent leaky integrate and fire network with 100 excitatory neurons with Hebbian plasticity. Our model was used to test whether the synaptic dynamics observed in the aged and young brain, could allow recovery from injury. To test general memory performance, we used an associative memory task, and tested the network under different conditions (i.e. before and after injury, in the aged and young brain).

## Stimulation Protocols

The model consisted of *N* excitatory neurons. Connections occurred with probability *P_con_*, and the strength of a connection from neuron *i* to neuron *j* was denoted by *W_ij_* (Fig.7, left). We aimed to train 3 “patterns”, which are 3 groups of highly connected neurons (Fig. 7, right). They were trained with *reps_train_* = 4 repeated presentation of external current with randomised order (Fig. 8A). These trained patterns are then used to test performance in a memory task. We used an associate memory task to test recall, by stimulating a percentage (60%) of the neurons in each pattern, with high external current, as in the training (pattern neurons = *N_pattern_*, test neurons = *N_test_*). Performance is then recorded using the error metric by counting the total amount of spiking of neurons that were not directly stimulated (Fig. 9 B). Performance would then be compared across different conditions in the baseline, and injury simulations. The total training time was *Sim_train_* = 1.8*s*, and the total testing time was *Sim_test_* = 2.61*s*. During training or testing, external current with rate of *r_Iextr_* or *r_Iexte_* Hz (train vs test), and of strength *I_extr_* or *I_exte_* pA (train vs test), which is equivalent to the sum of ∼5 or ∼15 (train vs test) input neurons, was activated sequentially for a period of *sim_train_* or *sim_test_*, with a gap of *g_train_* or *g_test_* between stimulus presentations. Neurons were targeted by each external stimulation with rate of *r_Iext_* Hz, and by noise with rate of *r_noise_*. As a result, during training, firing rate was on average 10 Hz and 26 Hz (EPBs vs TBs), comparable to firing frequencies in the somatosensory cortex (11 ± 0.82 Hz)^46,47^ with a maximum firing rate of 80 Hz and 148 Hz (EPBs vs TBs) during the stimulation. In testing, the average firing rate was 57 Hz and 50 Hz (EPBs vs TBs), and the maximum during the stimulation was 184 Hz and 165 Hz (EPBs vs TBs). While this is a high firing rate in comparison to most living neurons, it does not surpass the highest firing rates observed in the somatosensory cortex (185 *±*10 Hz).^46,48^

**Table 1.** Parameters for simulation.

| Parameters | Description | EPB | TB |
| --- | --- | --- | --- |
| $I_{ext_r}$ | Strength of external current stimulation in training | 3 pA | 3 pA |
| $I_{ext_e}$ | Strength of external current stimulation in testing | 10 pA | 10 pA |
| $r_{ext_r}$ | Rate of external current stimulation in training | 400 Hz | 500 Hz |
| $r_{ext_e}$ | Rate of external current stimulation in testing | 270 Hz | 270 Hz |
| $r_{noise}$ | Rate of noise stimulation | 100 Hz | 100 Hz |
| $N_{pattern}$ | Number of neurons in each pattern | 33 | 33 |
| $N_{test}$ | Number of test neurons | 13 | 13 |
| $Sim_{train}$ | Total training length | 1800 ms | 1200 ms |
| $Sim_{test}$ | Total testing length | 2610 ms | 2610 ms |
| $sim_{train}$ | Training external current stimulation length | 50 ms | 50 ms |
| $sim_{test}$ | Testing external current stimulation length | 270 ms | 270 ms |
| $g_{train}$ | Time between stimulations in training | 100 ms | 50 ms |
| $g_{test}$ | Time between stimulations in testing | 20 ms | 20 ms |
| $reps_{train}$ | Repetitions of stimulations in training | 4 | 4 |
| $reps_{test}$ | Repetitions of stimulations in testing | 3 | 3 |
| $dt$ | Time-step | 0.1 ms | 0.1 ms |

These parameters were fitted to induce the high connectivity in the patterns during training, to have enough stimulation to induce associative memory during testing, and also to ensure that firing rates remained at biological levels.

## Metrics

To calculate the performance of each model, we used the error metric. In order to compute the error, during the test we counted all the test neurons that fired during each test simulation for each pattern. Neurons that fired at the correct time (i.e. during the test current stimulation), were counted as True Positives (TP), and those that fired at other times were counted as False Positives (FP). We used TPs to compute the Error (E) metric:

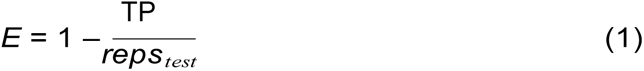

where *reps_test_*is the number of test stimulations per pattern.

## Injury Model

To model injury, we generated a weight matrix for each condition with parameters fitted (Fig.10, Table 2) using experimental data. We performed the following comparisons in our simulations: (1) Aged EBP-rich axons vs young EBP-rich axons (*n* = 50, repeats = 3), and (2) Aged EBP-rich axons vs young TB-rich axons (*n* = 50, repeats = 3, we fitted different biophysical values for TB-rich axons since they originate in L6, in contrast to EPB-rich axons which originate in L2/3/5 and thalamus). Since TB-rich axons regenerated more than EBP-rich axons in the young brain, we expected better recovery there. For each condition we generated a weight matrix for each of the steps (Fig. 10B). The steps include; baseline (i.e. pre-lesion), injury (i.e. boutons lost straight after injury), and post-lesion (i.e. re-wiring). At each step we performed an associate memory test, which is preceded by one repeat of a training simulation with Hebbian learning, aimed to mimic a more realistic dynamic environment as in the living brain. In more detail, we cut *Bloss_injury_* = 34% of connections from the trained weight matrices (Fig. 9A), by setting random connections to 0 in the weight matrix (calculated as average percentage of boutons lost via the injury). After injury, we update the weight matrix to simulate the results we observed in the live animal experiments (for example of weight matrix modification, see Fig. 10). In the aged EPB model, *Bgain_i_*_1_ = 18.8% connections are gained (Connections are added with weights randomly initialised between 0 to w_init_, and size is increased by *Bsize_i_*_1_ = 1.22 (stage 1). This is followed by *Bloss_i_*_2_ = 18.8% connections are lost (connections are removed by setting weights to 0), and size is decreased by *Bsize_i_*_2_ = 1.22 back to baseline levels (stage 2). In addition, the new connections from the previous step are increased by *BsizeNew_i_*_2_ = 2.41, as seen in the experimental results in this paper (stage 2).

- In the young EPB model, *Bgain_i_*_1_ = 2.5% connections are gained
- In the young TB model, *Bgain_i_*_1_ = 26% connections are gained, and an additional

**Table 2.**
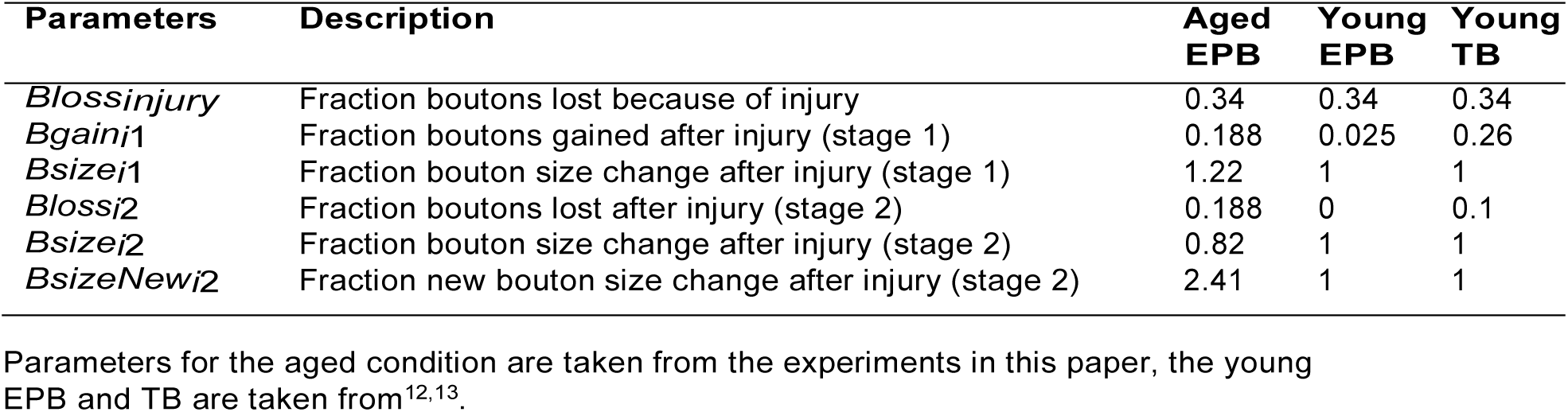
Parameters for injury model.

| Parameters | Description | Aged EPB | Young EPB | Young TB |
| --- | --- | --- | --- | --- |
| *Blossinjury* | Fraction boutons lost because of injury | 0.34 | 0.34 | 0.34 |
| *Bgaini1* | Fraction boutons gained after injury (stage 1) | 0.188 | 0.025 | 0.26 |
| *Bsizei1* | Fraction bouton size change after injury (stage 1) | 1.22 | 1 | 1 |
| *Blossi2* | Fraction boutons lost after injury (stage 2) | 0.188 | 0 | 0.1 |
| *Bsizei2* | Fraction bouton size change after injury (stage 2) | 0.82 | 1 | 1 |
| *BsizeNewi2* | Fraction new bouton size change after injury (stage 2) | 2.41 | 1 | 1 |

Parameters for the aged condition are taken from the experiments in this paper, the young EPB and TB are taken from<sup>11,12</sup>.

*Bloss_i_*_2_ = 1% of connections are lost post-lesion After each step (update to weight matrix), we run one repeat of training simulation (same parameters as in training, one repeat instead of 4, with Hebbian learning) to mimic dynamic conditions in the brain and passing of time, and then we perform an associate memory task (as described previously) to measure performance. We aimed to examine whether recovery from injury is possible with these given synaptic dynamics.

## Model Limitations and Assumptions

We use a simplified model, which is advantageous due to reduced number of parameters that need to be fitted, but there are a few limitations in how representative the model is of real neural networks:

- Integrate and fire neurons are relatively reductionist and do not represent full neuron dynamics, such as refractory period, adaptation, bursting, as they do not code memory of previous spikes, but just reset the voltage to the same value.^21^ These properties could be added to the model in different ways, but that would increase complexity and model parameters.
- Rate based Hebbian plasticity does not take into account pre and post pairing dynamics of action potentials in comparison to other models such as Spike-Timing-Dependent Plasticity (STDP).^21^ However, STDP learning rules can lead to pathological behaviour in balanced networks^49^, and can be difficult to set up well.
- We model only excitatory neurons, which is not fully representative of the cortex. Moresophisticated models with complex dynamics for a balanced network (excitatory and inhibitory neurons), have been developed^50^, but have a larger number of free parameters compared to our model. In addition, since we performed all our animal experiments on excitatory neurons, we decided it would be more appropriate to have a network of excitatory neurons, since we do not know how inhibitory neurons would behave following an injury.
- The time-scales in the model are in seconds, whereas the time-scales in animal experiments were in hours or days. Computationally, since simulating a model for such long periods would take an unrealistic amount of time, we reduce the time-scale to deal with the issue. Also, as we calculate TOR values as a ratio between 2 different simulations (with different parameters), we still fit to the desirable values as found experimentally.
- Since no experimental results were obtained for TB-rich axons in the aged brain, we were not able to run simulations for this condition. It is possible that since TB-rich axons have higher regeneration in the young brain, we would have also observed regeneration in the aged brain.
- We perform simulations for a relatively small number of neurons (*n* = 100). This can drastically reduce computational resources, and has worked well in our simulations.

In addition, we make several assumptions about the model:

- The turnover rate between connections of neurons was used to model ageing, as this is one of the key effects of ageing on structural plasticity.^9,10^
- To observe the effects on groups of connected neurons or populations of neurons, rather than single neurons, we set up our model as a group of randomly connected neurons. This setup is different to our animal experiments, where we image synapses on a single neuron and study their dynamics over time, and in response to injury. In our model each connection between two neurons is equivalent to a bouton or a synapse, as is measured in the animal experiments, i.e. a connection weight in our model is relative to the size or efficacy of a synapse that is imaged by two-photon microscopy.
- We assume different biophysical properties for EPB and TB-rich axons, as they originate at different areas in the cortex (L2/3/5 and the thalamus, and L6, respectively).^45^
- We assume that following injury re-wiring occurs most optimally-where connections are added to the patterns, and removed from outside the patterns. This however, might not be the case, so we perform a baseline where connections are added randomly. With random re-wiring we found similar but slightly worse recovery from injury than when using optimal re-wiring (see Appendix for results).

Overall, we tried to keep the model simple, to be able to isolate the effects of the synaptic dynamics, observed in our and other experiments on memory function. We validated our model by varying *dt* to ensure that the model works correctly (see *dt* experiment in Appendix). Finally, we attempted to use as many parameters as possible from biological experiments, for biological plausibility.^9,12,13^

## Membrane Potential Dynamics

The leaky integrate and fire model^51,52^ that we used is defined below:

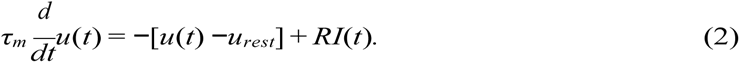

The internal current is updated at each time-step according to neuron firing:

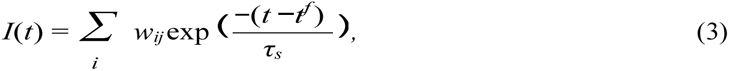

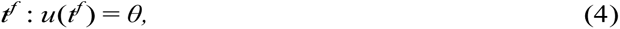

where *u*(*t*) is the time-varying membrane voltage, *u_rest_*is the resting membrane potential; *R* is the membrane resistance, *τ_m_* is the membrane time constant, *τ_s_* is the spike time constant, *I*(*t*) is the associated time-varying current, *w_ij_* is the synaptic strength between presynaptic neuron *i* and postsynaptic neuron *j*, *t^f^*is the *f^th^* firing time, and *θ* is the firing threshold voltage. A reset condition is also applied if the membrane potential goes over a threshold, *θ* (*−*38*mV*):

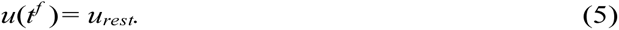

## Plasticity

We implemented a rate based Hebbian learning rule.^21,54^ A weight change, *dw_ij_/dt*, is applied depending on the firing rates of presynaptic and postsynaptic neurons (*ν_i_, ν_j_*, respectively):

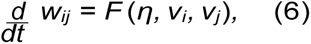

where *η* is the learning rate, and *F* is the learning function which is shown in the equation below and summarised in Table 4.

**Table 3.** Parameters for membrane neural dynamics.

| Parameters | Description | EPB | TB |
| --- | --- | --- | --- |
| $\tau_m$ | Membrane time-constant = RC | 28 ms | 28 ms |
| $U_{rest}$ | Reset voltage | -72 mV | -66.8 mV |
| $\theta$ | Threshold voltage | -38 mV | -40.2 mV |
| $R$ | Membrane resistance | 188 MΩ | 277 MΩ |

All parameters were taken from<sup>58</sup>

**Table 4.** Hebbian learning rule *F*.

| | $v_j < \langle v_j \rangle$ | $v_j > \langle v_j \rangle$ |
| --- | --- | --- |
| $v_i < \langle v_i \rangle$ | nothing | $\downarrow w_{ij}$ |
| $v_i > \langle v_i \rangle$ | $\downarrow w_{ij}$ | $\uparrow w_{ij}$ |

*w_ij_* is the synaptic strength between neurons *i, j*; *ν_i_, ν_j_*are presynpatic and postsynaptic firing rates, respectively; *(ν_i_)* and *(ν_j_)* are the thresholds for firing; *↑*, increase connection; *↓*, decrease connection.

The firing rate of each neuron *ν* is updated at each time-step:

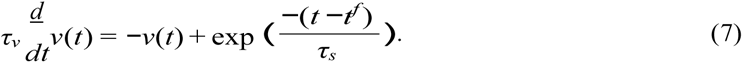

The weight update is similar to the covariance rule^54^ with a heaviside function:

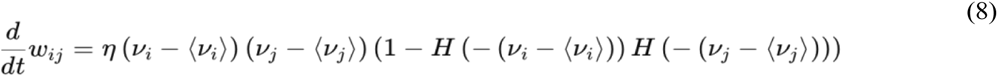

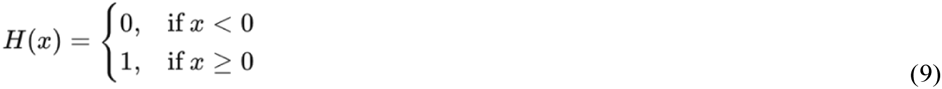

where *τ_ν_* is the time constant of low-pass filtered spike train, *H* is the Heaviside step function, *(ν)* is the average threshold for the firing rate for a neuron over time window *t_w_* (i.e. the firing rate threshold is updated as a running average over recent history), and *η* is the learning rate. The weight between two neurons, *w_ij_* is bounded so that it is kept between a lower and an upper bound,

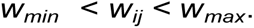

### Parameter Fitting

To use suitable parameters for EPB and TB models, we found the connectivity and the biophysical properties from the somatosensory cortex^53^ (Table 3). Since TB-rich axons originate from L6, and EPB-rich axons from L2/3/5 and thalamus, we used biophysical properties from these respective layers. We found that EPBs are known to be bigger and more stable [19], so we gave them a higher weight boundary (which corresponds to their size) than TBs (0.64 and 1.28 weight boundaries for TBs and EPBs, respectively, Table 6). Another property we tried to keep within physiological range is the firing rate, which has been found to be on average 11 ± 0.82 Hz^46,47^ and maximally 185 ± 10 Hz for neurons in the somatosensory cortex.^46,48^ For the aged and young models, we fitted the TOR (same calculation as in experiments) of young (EPBs and TBs) and aged (EPBs) synapses.^9,12,13^ TOR is computed based on gains and losses-i.e. a gain and loss is when the weight goes over or under *θ_synaptic_* threshold. We computed TOR at each simulation for different learning rates (*β*). We then made sure that the TOR of each group (e.g. young adult EPB, and aged EPB) is in the correct ratio to each other (within an error of 0.01). For example, in the aged EPB group the TOR from experimental methods is 0.15, and the TOR in the young EPB group is 0.08. For each pair of learning rates we checked that TOR in the aged group is 1.85X higher than in the young group. We did the same for the other groups (aged EPB vs young TB; young EPB vs young TB). The parameters used in the model for the injury model are summarised in Table 2. We first extracted parameters from existing publications^12,13^ where we computed that following injury EPB-rich axons regenerate, and gain boutons equal to the number of boutons lost divided by 13.7. This was computed by looking at the ratio of number of boutons lost from injury, to number of boutons gained from the regeneration. In a similar manner we also computed this ratio for TB-rich axons, and found that the number of boutons gained following injury is equal to the number of boutons lost divided by 1.29 (i.e. there is more regeneration in TB-rich axons). In TB-rich axons there was also a pruning mechanism following injury, where 0.1 of the boutons were lost. We then did a comprehensive analysis of the bouton dynamics shown in this paper to use in the model. We split the simulation of the aged EPB axons to two stages (see Table 2 for a complete breakdown for each stage); (1) first stage for the 6 hour time-point where we observed an increase in number and size of boutons, and (2) second stage for these return back to baseline. More specifically, we found that 6 hours following injury that is a 1.22 increase in size of boutons, and 1.188 increase in number of boutons (which is calculated based on the boutons that are currently present). The new boutons that are gained were found to increase their size by 2.41 before injury. In the results section, for readability, we report only the final result of the second stage.

**Table 5.** Parameters for synaptic plasticity.

| Parameters | Description | Values |
| --- | --- | --- |
| $\beta$ | Hebbian learning rate | $0.01 \leq \beta \leq 0.05$ pA |
| $\tau_v$ | Time constant of low-pass filtered spike train | 20 ms |
| $\tau_s$ | Spike time-constant | 0.01 ms |
| $t_w$ | Time-window for firing rate | 10 ms |
| $\theta_{synaptic}$ | Weight threshold for synapse gain or loss | 0.4 mV |

**Table 6.** Parameters for network graph.

| Parameters | Description | EPB | TB |
| --- | --- | --- | --- |
| $w_{max}$ | Upper weight boundary in learning | 1.28 mV<sup>29,58</sup> | 0.64 mV<sup>58</sup> |
| $w_{min}$ | Lower weight boundary in learning | 0 mV | 0 mV |
| $w_{init}$ | Weights are initialised between 0 to this value | 0.64 mV | 0.64 mV |
| $P_{con}$ | Probability of connections between neurons | 0.23<sup>58</sup> | 0.23<sup>58</sup> |
| $N$ | Number of neurons in the network | 100 | 100 |

## Appendix

### Additional Injury Experiments

We performed an additional baseline experiment, to explore whether if the re-wiring following injury in non-optimal, i.e. when boutons are gained following injury it is added randomly to the entire weight matrix, and similarly when boutons are removed, they are removed randomly from the entire weight matrix (Fig. A1).

**Figure A1.**
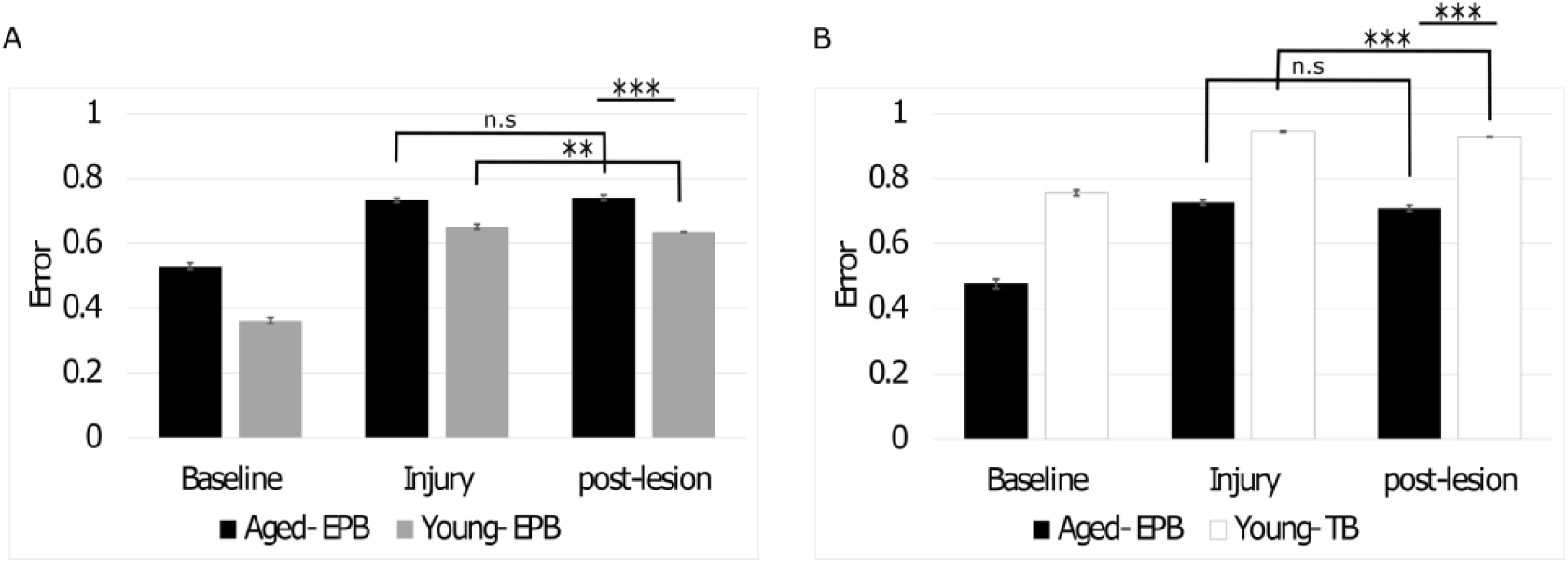
Results for injury simulation with random wiring. **A**, No difference between injury and post-lesion in the aged EPB condition, recovery of 0.02 (*p <* 0.01) in the young EPB condition. **B**, No difference between injury and post-lesion in the aged EPB condition, recovery of 0.02 (*p <* 0.001) in the young TB condition. Error bar, SEM.

As expected, recovery is less in this case, but we still observe significant recovery in the young EPB (*p <* 0.01), young TB (*p <* 0.001) conditions, and non-significant recovery in aged EPB (*p* = 0.09) condition. It is likely that the brain performs re-wiring in a more optimal manner, if its aim is to recover its synaptic dynamics. The true re-wiring mechanism, might be between the random re-wiring to optimal re-wiring that we tested in these simulations.

### Changing *dt* Experiments

We validated whether our model works correctly when changing *dt* by a small value (*dt* = 0.1, 0.095). We compared the training and testing simulation graphs, weight matrices and the firing rates, and found that they were comparable (repeats = 5). During training (Fig A2, left) the average firing rate was 7 ± 0.4Hz and 9.5 ± 0.9Hz (*dt* = 0.095, and 0.1, respectively). The firing rates during testing was also comparable (Fig A2, middle). Additionally on average (repeats = 5), the sum of the weights was nearly identical (Fig A2, right) 0.1772 ± 0.01 and 0.1778 ± 0.01 (*dt* = 0.095, and 0.1, respectively). This demonstrates that our model is correctly normalising the simulations by *dt*. Changing *dt* by a large amount is likely to change the results, as the model accuracy changes (i.e. the smaller the change in time *dt*, the higher the accuracy) with significant change in *dt*.

**Figure A2.**
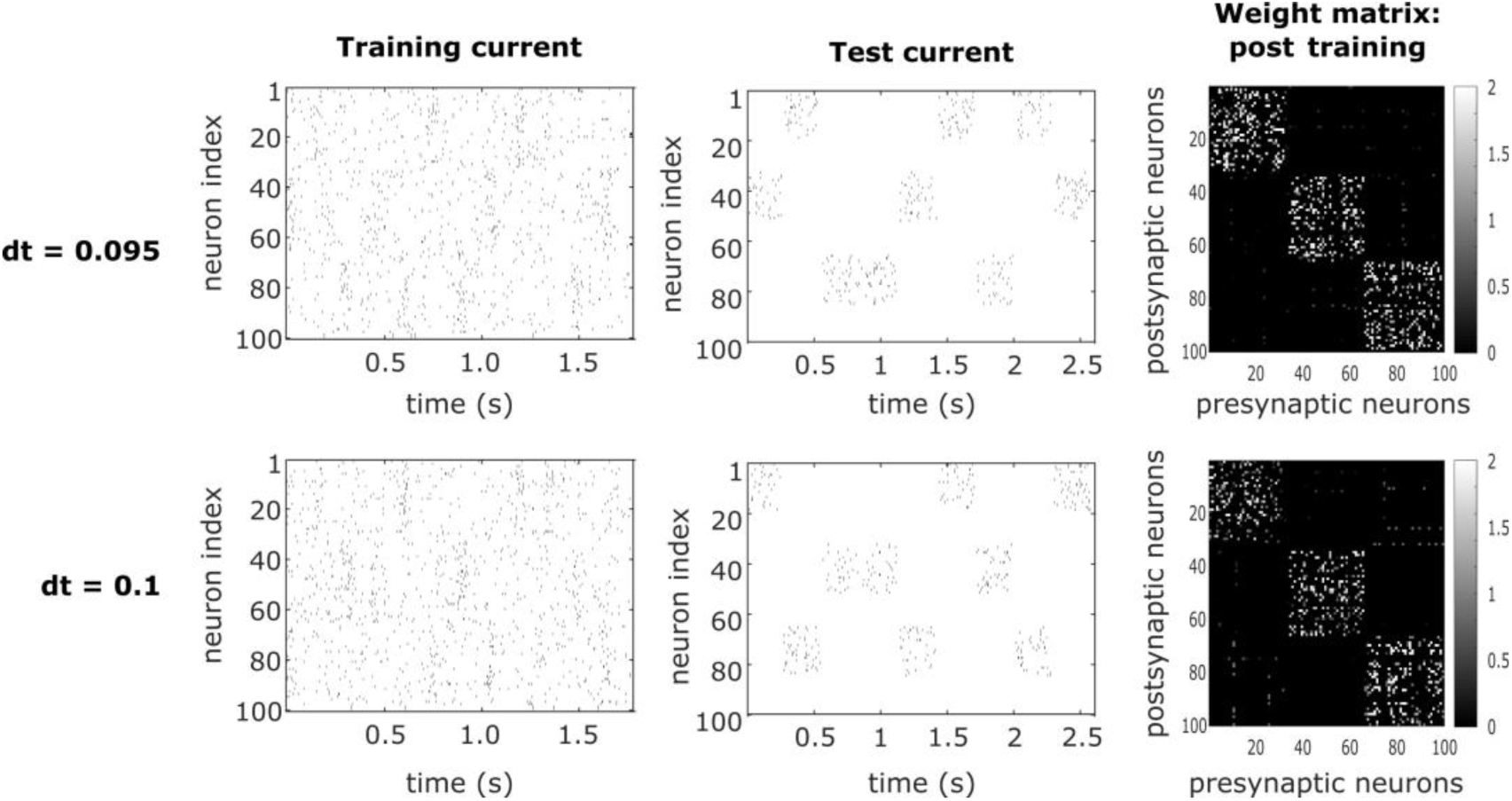
Plots for simulations with varying *dt*s. Example training and testing current plots, and weight matrices for each simulation, with different *dt*s (0.095 and 0.1).

